# The partial duplication of an E3-ligase gene in *Triticeae* species mediates resistance to powdery mildew fungi

**DOI:** 10.1101/190728

**Authors:** Jeyaraman Rajaraman, Dimitar Douchkov, Stefanie Lück, Götz Hensel, Daniela Nowara, Maria Pogoda, Twan Rutten, Tobias Meitzel, Caroline Höfle, Ralph Hückelhoven, Jörn Klinkenberg, Marco Trujillo, Eva Bauer, Thomas Schmutzer, Axel Himmelbach, Martin Mascher, Barbara Lazzari, Nils Stein, Jochen Kumlehn, Patrick Schweizer

**Affiliations:** Leibniz Institut für Pflanzengenetik und Kulturpflanzenforschung (IPK Gatersleben), Corrensstrasse 3, D-06466 Stadt Seeland, Germany; Technische Universität München, Emil-Ramann-Straße 2, D-85354 Freising, Germany; Leibniz Institut für Pflanzenbiochemie, Weinberg 3, D-06120 Halle (Saale), Germany; Technische Universität München, Liesel-Beckmann-Straße 2, D-85354 Freising, Germany; Parco Technologico Padano, Via Einstein, Loc. Cascina Codazza, 26900, Lodi, Italy

**Keywords:** Hordeum vulgare, Blumeria graminis, Triticeae, Plant U-box protein, Armadillo-repeat, Partial gene duplication

## Abstract

In plant-pathogen interactions, components of the plant ubiquitination machinery are preferred targets of pathogen-encoded effectors suppressing defense responses or co-opting host cellular functions for accommodation. Here, we employed transient and stable gene silencing-and over-expression systems in *Hordeum vulgare* (barley) to study the function of *HvARM1* (for *H. vulgare Armadillo 1*), a partial gene duplicate of the U-box/armadillo-repeat E3 ligase Hv*PUB15* (for *H. vulgare Plant U-Box 15*). The partial *ARM1* gene was derived from an ancient gene-duplication event in a common ancestor of the *Triticeae* tribe of grasses comprising the major crop species *H. vulgare*, *Triticum aestivum* and *Secale cereale*. The barley gene *HvARM1* contributed to quantitative host as well as nonhost resistance to the biotrophic powdery mildew fungus *Blumeria graminis*, and allelic variants were found to be associated with powdery mildew-disease severity. Both HvPUB15 and HvARM1 proteins interacted in yeast and plant cells with the susceptibility-related, plastid-localized barley homologs of THF1 (for Thylakoid formation 1) and of ClpS1 (for Clp-protease adaptor S1) of *Arabidopsis thaliana*. The results suggest a neo-functionalization *HvARM1* to increase resistance against powdery mildew and provide a link to plastid function in susceptibility to biotrophic pathogen attack.

## INTRODUCTION

Plants response to pathogen attack by the activation of their innate-immunity system, which is triggered by the perception of pathogen-associated molecular patterns (PAMPs). On the other hand, successful plant pathogens manipulate their hosts by complex arsenals of secreted effector proteins to co-opt cellular host functions for accommodation, nutritional exploitation, or for suppression of immunity. A growing number of these were found to target components of the plant ubiquitination machinery including plant U-box E3 ligases (PUB’s) (Abramovitch *et al.*, 2006, Angot *et al.*, 2006, Bos *et al.*, 2010, Groll *et al.*, 2008, Nomura *et al.*, 2011, Rosebrock *et al.*, 2007, Spallek *et al.*, 2009). The covalent attachment of single ubiquitin moieties or polyubiquitin chains to lysine residues of eukaryotic protein substrates can have diverse effects on their fate. Ubiquitination most commonly results in the recognition and degradation of tagged proteins by the 26S proteasome but also mediates endosomal sorting into cellular compartments such as the lysosome or the plant vacuole, or contributes to DNA damage responses (Trujillo & Shirasu, 2010, Vierstra, 2009). The substrate specificity during ubiquitination is determined by the E3 ubiquitin ligases which can be sub-divided into three categories namely HECT, RING/U-box type, and cullin-RING ligases. These proteins mediate ubiquitin ligation in concert with the highly conserved ubiquitin-activating enzyme (E1) and ubiquitin-conjugating enzymes (E2). Due to their central cellular function, components of the ubiquitination system represent central cellular hubs of protein regulation involved in all aspects of plant life. As such, beneficial or parasitic organisms may utilize the ubiquitination machinery (Spallek *et al.*, 2009) to establish susceptible interactions. On the other hand higher plants exploit ubiquitin-mediated degradation of negative protein regulators of stress-hormone signaling for the initiation of PAMP-triggered immunity (PTI) that appears often to underlie quantitative host resistance (QR) or nonhost resistance (NHR) (Sadanandom *et al.*, 2012, Schweizer, 2007, Trujillo & Shirasu, 2010).

NHR protects plants from the vast majority of attacks by pathogens that have adapted during co-evolution to different, more or less closely-related plant species (Schulze-Lefert & Panstruga, 2011). Another broadly acting form of resistance against pathogens is QR, which is usually determined by several QTL, as revealed by crossing more resistant genotypes with susceptible ones, and is generally looked upon as a manifestation of PTI. In contrast to ETI it does not confer complete protection but may be more durable in the field. Cultivated barley (*Hordeum vulgare* ssp. *vulgare*) is a nonhost to the non-adapted wheat powdery mildew fungus *Blumeria graminis* f.sp. *tritici* (*Bgt*) but a host of the powdery mildew fungus *B. graminis* f.sp. *hordei* (*Bgh*) causing up to 30% yield loss in the absence of genetic or chemical control of the disease (Oerke, 2006, Panstruga & Schulze-Lefert, 2002). The epidemic spread of *B. graminis* is caused by the asexual propagation of the fungus, with a generation time of 5-7 days and massive production of conidiospores (Figure 1). The interaction between different barley genotypes and *Bgh* isolates represents a well-studied model system for a fungal disease caused by an obligate biotrophic pathogen, and a growing number of host-response factors for defense or disease establishment have been identified (Collinge, 2009, Huckelhoven, 2007, Huckelhoven & Panstruga, 2011). The genome of *Bgh* was found to encode over 500 candidate secreted effector proteins (Pedersen *et al.*, 2012). As described in plant-pathogenic *Pseudomonas* sp. bacteria or in filamentous *Oomycete* pathogens, *Bgh* effectors are likely to target host ubiquitination components, too. However, up to present this remains speculative based on conceptual considerations and extrapolation. In a phenotype-driven, transient RNA interference (RNAi) screen for the discovery of *Rnr* (for *Required for nonhost resistance*) genes to the non-adapted wheat powdery mildew fungus *B.graminis* f.sp. *tritici* (*Bgt*), we tested over 631 barley genes, which were mostly associated with up-regulated transcripts in *Bgt*-attacked barley leaf epidermis (Douchkov et al., 2014). Reduced NHR was reflected by an increased percentage of transformed epidermal cells containing *Bgt* haustoria. Out of 46 RNAi target genes that fulfilled the selection criteria during the first screening round of transient-induced gene silencing (TIGS), 10 final *Rnr* gene candidates significantly enhanced nonhost susceptibility upon silencing.

**Figure 1:**
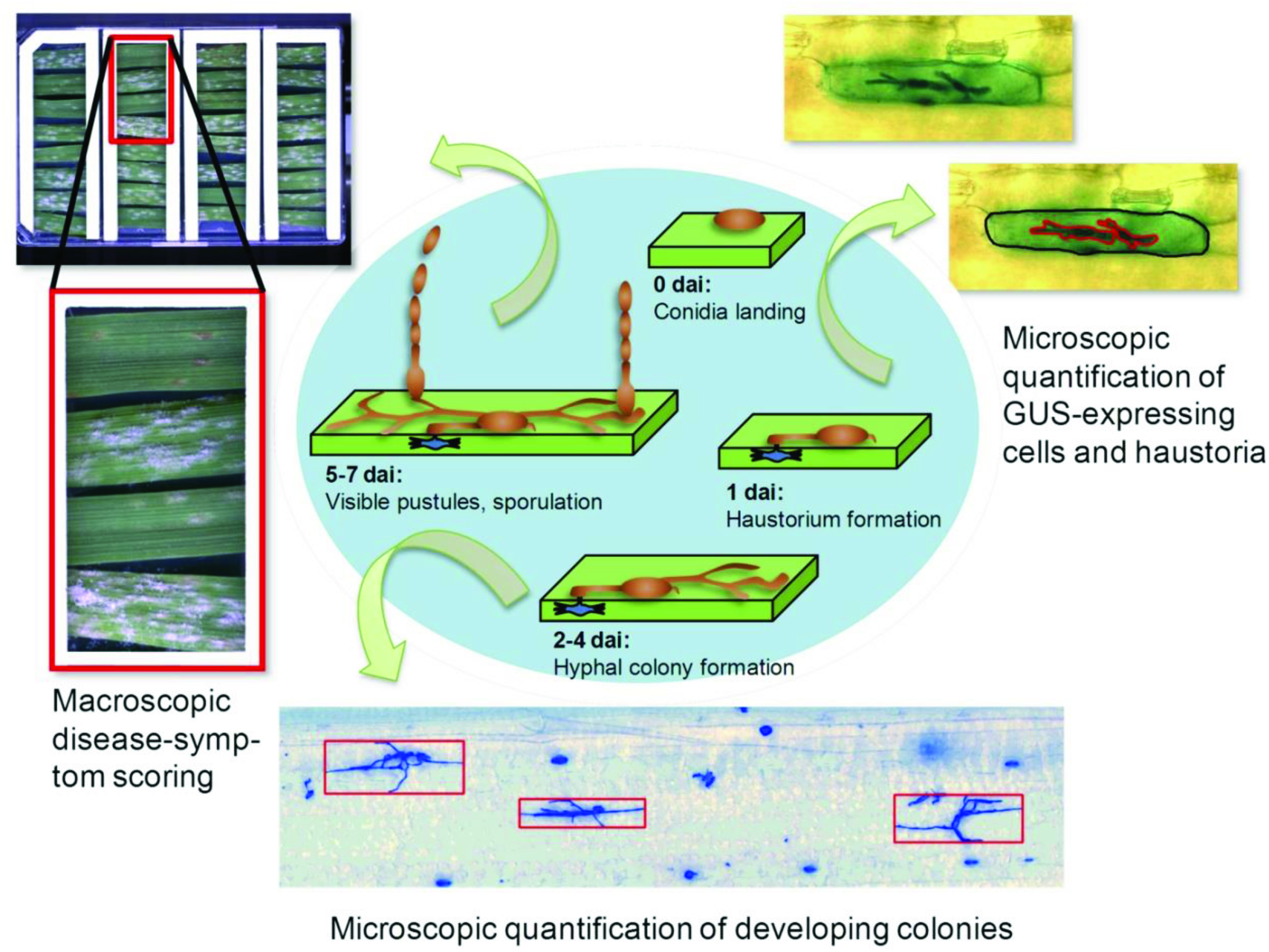
Asexual life cycle of *Blumeria graminins* and the assessment of fungal development in barley (*Hordeum vulgare*) and wheat (*Triticum aestivum*). For transient-induced gene silencing (TIGS) and for transient over-expression (OEX), initial haustorium (feeding cell) formation in transformed, GUS-expressing epidermal cells of barley or wheat is completed 1 day after inoculation (dai) and was assessed under the microscope 40 h after inoculation. For the characterization of transgenic plants, early developing colonies were stained, counted and normalized to the analysed leaf area at 2 dai. The asexual life cycle is completed by the appearance of macroscopiclaly visible, sporulating colonies (pustules) 5-7 dai, which can be estimated or quantified as percentage of pustule-covered leaf area.

Here we present structural and functional data to validate *Rnr5* encoding Hv*ARM1* with homology to PUBs. Besides its possible role in NHR that allowed discovery in the NHR RNAi screening, functional analysis suggested *Rnr5* as an important factor of QR against the adapted *Bgh* fungus.

## MATERIALS AND METHODS

A more detailed description of materials and methods used in this study is provided in supporting Information.

### Plant and fungal material

For the TIGS and transient overexpression experiments 7-day-old seedlings of spring barley cv. Maythorpe and Golden Promise were used, respectively. As exception and for better comparability to transgenic plants, TIGS experiments for *HvPUB15* and *HvARM1* were also done in cv. G. Promise. Stable transgenic barley plants of cv. G. Promise were generated as described (Hensel *et al.*, 2008). Bombarded leaf segments or transgenic plants were inoculated with Swiss *Bgt* field isolate FAL 92315, or Swiss *Bgh* field isolate CH4.8 throughout the study.

### Exome capture sequencing

Genomic DNA was extracted from barley leaf material from a single plant for each accession and used for the hybridization with the barley SeqCap Ez oligo pool (Design Name: 120426_Barley_BEC_D04, (Mascher *et al.*, 2013). Quality-trimmed reads were mapped to the reference genome (http://webblast.ipk-gatersleben.de/barley_ibsc/downloads/) with BWA version 0.7.5a using the mem algorithm with default parameters (Li & Durbin, 2009) and retaining only properly paired reads. Variant calling and realignment around indels were performed with GATK, version 2.7.4 (https://www.broadinstitute.org/gatk/"). Variant calls were filtered for high quality and ≥ 80% of samples being represented at each locus, and a dataset of 449585 SNPs was produced, suitable for association-genetic analysis of the two genes under investigation (full information about genome wide variants from this dataset will be published elsewhere).

### Association genetic analysis

Association of SNP and gene-haplotypes (marker) of *HvARM1* and *HvPUB15* with the severity of *Bgh* infection (trait) was calculated based on genetic and phenotypic data of two diverse collections of cultivated barley (*H. vulgare* ssp. *vulgare). Bgh* infection values were determined in a detached leaf assay using second leaves of approximately 12-day-old seedlings, as described (Spies et al., 2012). First, a worldwide collection of 76 landraces (WHEALBI_LRC) was inoculated either with isolate JKI-75 or JKI-242 that exhibit a complex and complementing virulence spectra (Šurlan-Momirović *et al.*, 2016). Second, a worldwide collection of 127 cultivars (WHEALBI_CULT) was inoculated with the same 2 *Bgh* isolates. Both populations consisted of single seed-derived lines, and an average of 5 parallel plants per line was used in each inoculation assay. For passport data of all lines see supplemental Table S1. Seven days after inoculation, disease was scored by estimating the percentage of leaf area covered by fungal mycelium. Because disease scores were variable between different inoculation experiments they were normalized to internal standards cv. Roland or Morex, as indicated. Phenotypic data of all isolate-genotype combinations are based on 2 independent inoculation series. SNP calls were derived from exome capture re-sequencing, and haplotypes were calculated based on the combination of SNP calls per gene. SNP-trait and haplotype-trait associations were calculated in TASSEL v4.1 using a mixed linear model with kinship as random effect. Marker data for kinship calculations were derived from 4032 polymorphic “Genotyping by Sequencing” (GBS) markers. Marker-trait associations were assumed significant if the Holm’s-corrected p value was <0.05 (number or SNP or haplotypes/gene = number of tests).

### TIGS and transient over-expression

TIGS constructs were generated and transferred by particle bombardment into leaf epidermal cells of 7-day-old barley seedlings as described (Douchkov *et al.*, 2005). Leaf segments were inoculated three days after the bombardment with *Bgh* at a density of 140-180 conidia mm^−2^. Transformed GUS-stained epidermal cells as well as haustoria-containing transformed (susceptible) cells were counted 48 hours after inoculation, and TIGS effects on the susceptibility index (SI) were statistically analyzed (Spies *et al.*, 2012).

For transient overexpression, a Hv*ARM1*-containing BAC clone HVVMRXALLhA0581d24 (Acc. Nr. KM979563) was bombarded into leaf segments of barley cv. Maythorpe or wheat cv. Kanzler, followed by challenge inoculation with the corresponding adapted pathogen *Bgh* or *Bgt* 4 hours after the bombardment and microscopic assessment of SI 48 hours after inoculation. For verification of transgene effects, Hv*ARM1* was excised from a subclone of the above-mentioned BAC as StuI/SphI fragment and inserted into SmaI/SphI sites of pIPKTA09. For transient overexpression of candidate genes, full-coding sequences were PCR amplified from cDNA and inserted as *XbaI* fragment into to the multiple cloning site of pIPKTA09 (Zimmermann *et al.*, 2006). The resulting sequence-verified constructs were bombarded into barley as described for BAC clones. For PCR primers used in this study see supporting Table S6.

For the Thf1:YFP and ClpS1:YFP degradation assay (Figure 7), four µg of respective plasmid DNA plus pUbiGUS (Douchkov *et al.*, 2005) were co-bombarded into 7-day-old barley cv. G. Promise. The numbers of YFP-fluorescing cells with plastid-localized signals were counted 24 hours after particle bombardment, followed by GUS-staining (Douchkov *et al.*, 2005). The numbers of GUS-expressing cells were used for normalization of the YFP signal.

**Figure 7:**
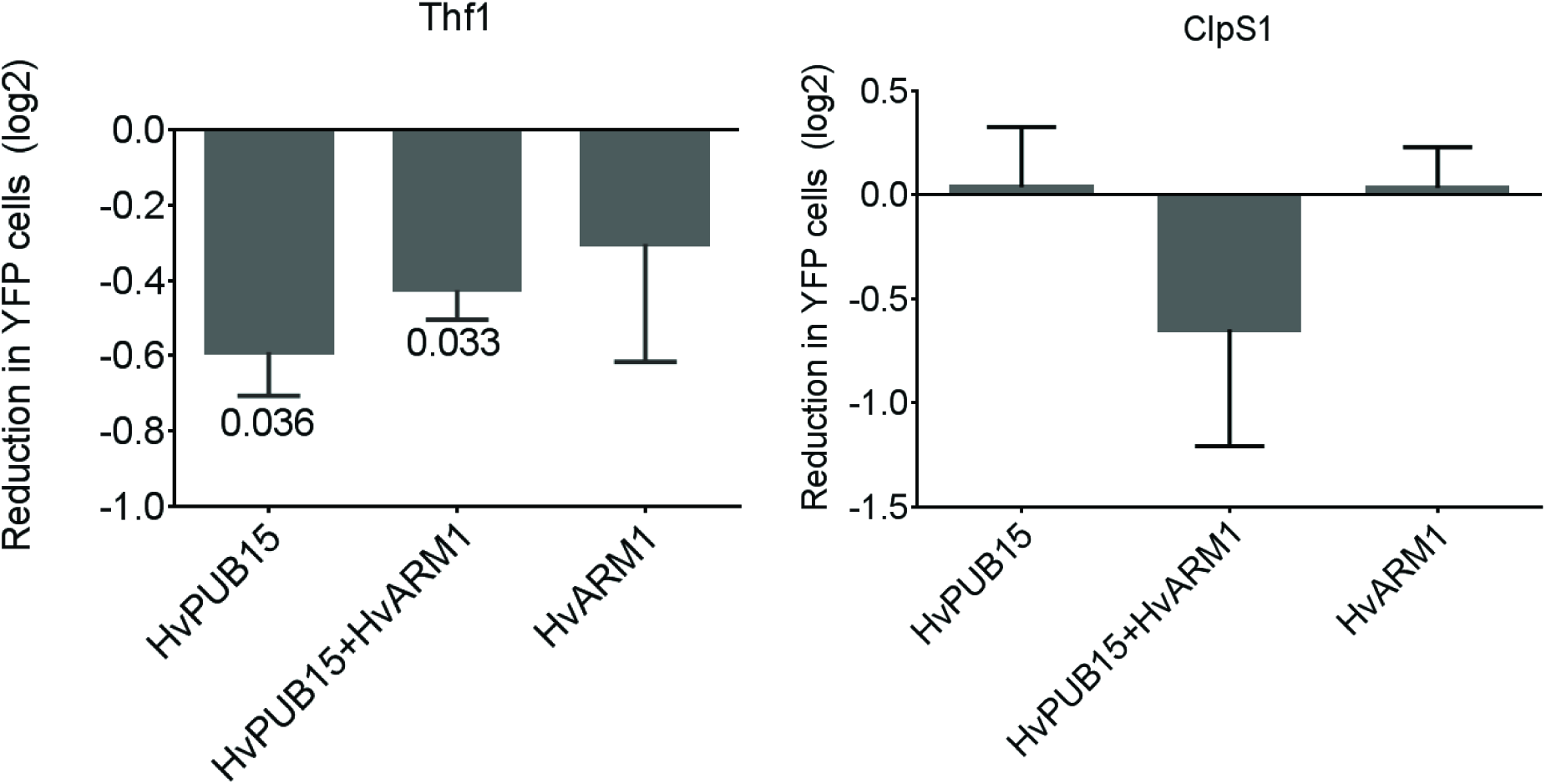
Degradation of HvThf1:YFP by over-expression of HvPUB15. The numbers of HvThf1:YFP-or HvClpS1:YFP-expressing epidermal cells was determined in transient co-expression experiments in presence or absence of HvPUB15 or HvARM1. Data represents log_2_-transformed values normalized to the empty-vector control. Bars indicates mean ± SEM of 3-4 independent bombardment series; P values for the null hypothesis are indicated (one sample t-test, two tailed).

### Inoculation and evaluation of transgenic plants

Phenotypic evaluation of *Bgh* and *Bgt* interactions was done microscopically on second, detached leaves of 12-14 day-old plants placed on phytoagar plates (23,2 cm x 23,2 cm) inoculated at a spore density of 30-40 conidia mm^−2^. Inoculated leaf segments were incubated for 48 hours (*Bgh*) or 72 hours (*Bgt*) followed by staining with Coomassie brilliant blue R 250 (Schweizer *et al.*, 1993). The number of growing colonies/leaf area was counted under a standard bright field microscope at 100 x magnification.

### Yeast-two-hybrid experiments

Yeast-two-hybrid screening was performed according to the Yeast Handbook and manual of Matchmaker™ library-construction and -screening kits (Takara/Clontech Laboratories, Saint-Germain-en-Laye, France). Full length coding sequence of Hv*ARM1* (1-442 AA) was used to screen a library of 7 × 10^6^ mating events according to (Hoefle *et al.*, 2011). For targeted Y2H assays, the coding region (1-831 AA) of HvPUB15 was used to test positive prey clones of the HvARM1 screening.

### Bimolecular fluorescence complementation and co-immunoprecipitation

For bimolecular fluorescence complementation (BiFC) of HvARM1 and HvPUB15 proteins with potential plastid interactors, *Nicotiana benaminiana* plants were grown and agro-infiltrated as described in detail in supporting Information Methods S1. For BiFC with HvThf1 and HvClpS1, the wild-type full-length sequences of *HvPUB15* or *HvARM1*, U-box mutants of *HvPUB15*, the ARM-domain (351 to 831 AA) only of *HvPUB15*, or *HvThf1* without N-terminal plastid import signal (-SP) were cloned into 35S∷^GW^VYNE-pBar and 35S∷^GW^VYCE-pBar GATEWAY destination vectors containing the N-and C-terminal split parts of the enhanced YFP protein *Venus*, respectively (Thormahlen *et al.*, 2015). BiFC constructs were transiently co-expressed by infiltration of *Agrobacterium tumefaciens* transformed with the corresponding binary vectors, and examined by confocal laser-scanning microscopy (CLSM) 48 hours after infiltration. For the development of U-box mutants, DNA fragment between 709-739 bp (from ATG) on the U-box domain of HvPUB15 was excised using BsaXI and replaced by ligating synthetic oligos carrying the respective U-box mutation.

For co-immunoprecipitation (Co-IP), YFP-tagged HvARM1 and HvPUB15^ARM^ under the control of 35S promoter were generated by cloning the full-coding sequence of Hv*ARM1* (1-442 AA) or the ARM-repeat region of Hv*PUB15* (351-831 AA) into pEARLEYGATE104 (Earley et al., 2006). cMyc-Tagged HvThf1 (1-294 AA) and Hv*ClpS1* (1-161 AA) under the control of the CaMV 35S promoter were generated by cloning into pGWB418 (Nakagawa et al., 2007). Mesophyll-protoplast transformation and co-immunoprecipitation was done as described (Stegmann *et al.*, 2012).

### Subcellular localization of fluorescent proteins

For subcellular localization, full-length sequences of Hv*PUB15*, Hv*ARM1*, HvThf1 and Hv*ClpS1* were N-and C-terminally fused in-frame to YFP in pIPKTA48 and pIPKTA49 vectors (supporting Information Figures S9 and S10). Resulting YFP-fusion constructs were transiently expressed in 7-day-old barley leaf segments by particle bombardment and examined after 12-24 hours of incubation with or without *B. graminis* inoculation using confocal laser-scanning microscopy (CLSM).

## RESULTS

### Origin and evolution of HvARM1

Transient single-cell silencing of *Rnr5* significantly reduced NHR of barley to *Bgt* (Douchkov *et al.*, 2014). BlastX analysis revealed homology of *Rnr5* to plant U-box type E3 ligases (PUBs) with an armadillo-(ARM) repeat as second conserved domain (Azevedo *et al.*, 2001, Mudgil *et al.*, 2004). Although *Rnr5* was most closely related to *OsPUB15* in rice (Zeng *et al.*, 2008) it does not appear to be the barley ortholog because the encoded protein of 442 amino acids is considerably shorter than a regular PUB and contains only the C-terminal ARM-repeat region as conserved domain (Figure 2a and supporting Figure S1). The barley genome also contains a gene for a full-length PUB protein of 831 amino acids with highest similarity to *OsPUB15* that was therefore named *HvPUB15*, whereas *Rnr5* was designated as *HvARM1* (Table 1). Protein similarity between HvPUB15 and HvARM1 starts at position L398 and L9 of HvPUB15 and HvARM1, respectively, between the conserved U-box and ARM-repeat regions of HvPUB15 (supporting Figure S2). Sequence similarity between the two genes extends upstream from the *HvARM1* initiating codon spanning the first intron of *HvARM1*, which corresponds to exon 3 sequence of *HvPUB15*, until it abruptly ends within the U-box sequence of Hv*PUB15*. Further upstream sequence inside *HvARM1* intron 1 as well as the non-translated exon1 sequence did not exhibit significant similarity to any annotated gene or repetitive DNA element in the barley genome. These results strongly suggest that *HvARM1* originated as a partial gene duplicate of *HvPUB15*. An 8-bp deletion downstream from the first five N-terminal amino acids of *HvARM1* restored the initial reading frame because its initiating ATG corresponds to an out-of-frame codon of Hv*PUB15* (Figure 2b and c). Whole-genome shotgun (WGS) sequences of three additional species of the *Triticeae* tribe of grasses, the wild wheat species *Aegilops tauschii* and *Triticum urartu* plus rye (*Secale cereale*), also revealed the presence of *HvARM1*-like genes suggesting a monophyletic origin of the partial gene-duplication event in a common *Triticeae* ancestor dating back at least 12 M years (Figure 2d). Protein-sequence conservation among the four species was found to be high in both the U-box containing N-terminal-and the ARM-repeat containing C-terminal part of PUB15, the existing polymorphisms being in agreement with phylogenetic species distances. By contrast, sequence conservation was reduced among ARM1 proteins, most clearly evident when comparing the two wild wheat species. To address the question if different evolutionary constrains act on *PUB15* and *ARM1* genes we searched both orthologous gene groups for footprints of purifying or diversifying selection by calculating Ka/Ks ratios in windows of 40 amino acids within the overlapping parts of both proteins. Because a Ka/Ks ratio of 1 indicates neutral selection, we tested mean Ka/Ks of all six possible pairwise sequence comparisons between the four species for significant differences from the null-hypothetical value “1”. As shown in Figure 2e, both genes are subjected to purifying selection at the very N-terminus of ARM1 and within the ARM-repeat region. Selection was neutral in ARM1 outside these regions whereas PUB15 sequences remained under purifying selection along the entire ARM1-overlapping part of the gene. This suggests that the function of ARM1 proteins is restricted to the binding of one or a few protein ligand(s) via its ARM repeats whereas structural constrains on full-length E3-ligases that have to bind to substrate proteins and mediate the interaction with the highly conserved UBC domain of E2’s are probably more stringent.

**Figure 2:**
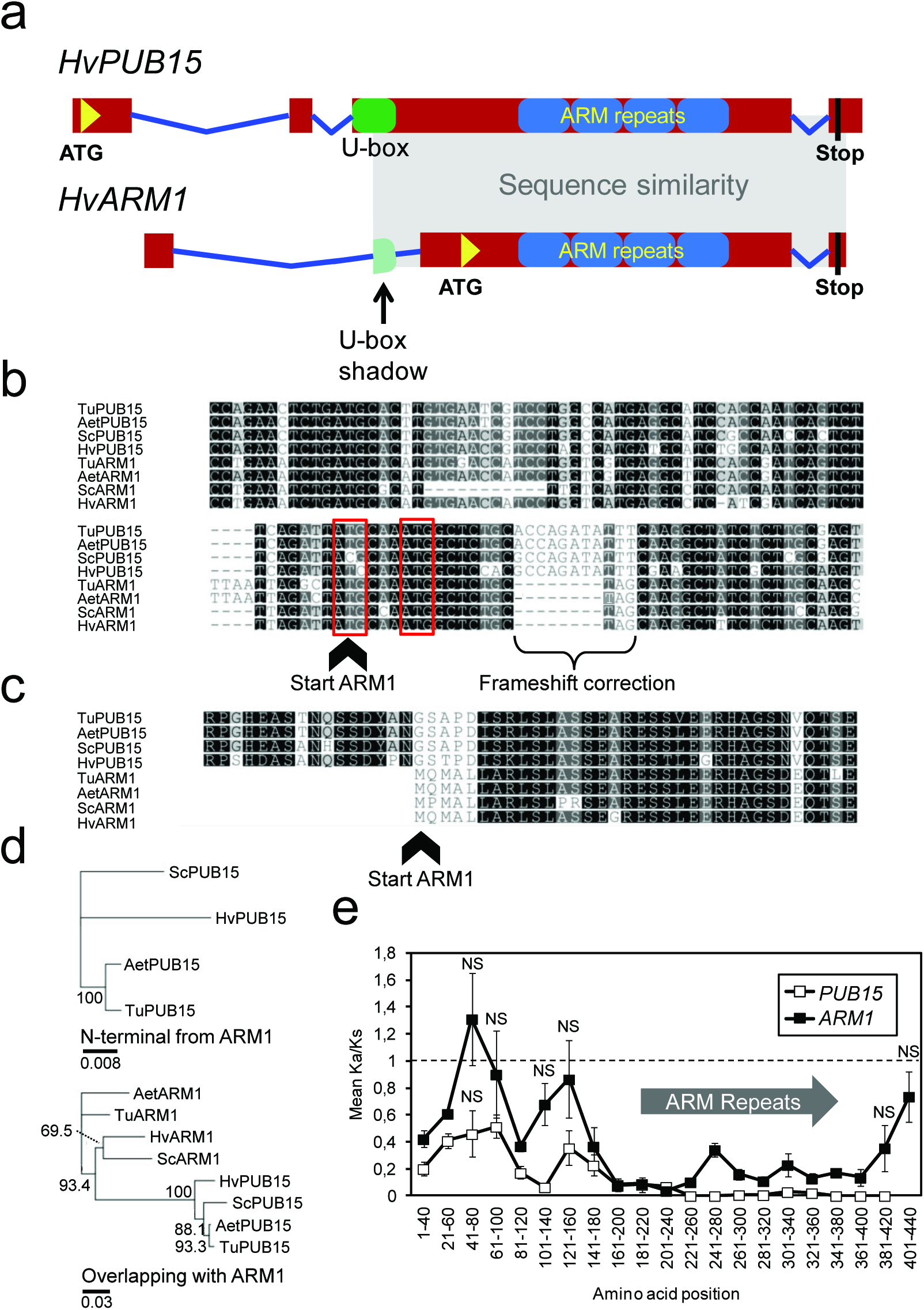
A partial duplication of the E3 ligase gene *PUB15* in *Triticeae* species gave rise to *ARM1*. **(a)**Schematic view of the genomic structure of HvPUB15 and its partial duplicate HvARM1. Red boxes and blue lines represent exon and intron sequences, respectively. The region of high sequence homology is indicated by light gray shading. **(b)**DNA-sequence alignment around the translational start of *ARM1* from *Triticeae* species. The two proposed translation start sites of *ARM1* are inside the red frames. Percent sequence identity per nucleotide is indicated by grey shading (black, 100% identity). Acc-Nr: *AetARM1*, XM_020296982; *HvARM1*, AK371875; *ScARM1*, KM881628; *TuARM1*, AOTI010733077; *AetPUB15*, XM_020311842; *HvPUB15*, AK361754; *ScPUB15*, KM881629; *TuPUB15*, AOTI010384858. (c) Protein sequence alignment of PUB15 and ARM1 at the N-terminus of ARM1. Acc. Nr: AetARM1, EMT05611; HvARM1, BAK03073; ScARM1, KM881628; TuARM1, EMS55038; AetPUB15, EMT16948; HvPUB15, BAJ92958; ScPUB15, KM881629; TuPUB15, EMS61710.**(d)**Phylogenetic trees of PUB15 protein sequences based on alignment from the N-terminus to the start of the overlapping part with ARM1, and of both proteins based on alignment of overlapping PUB15 and ARM1 sequences. A neighbour-joining tree without pre-determined outgroup was calculated, and bootstrap values (in percent) based on 1000 re-iterations plus tree depth (in changes per amino-acid position) are indicated by numbers and scale bar, respectively. **(e)**Conservative selection at the armadillo-repeat domain of *ARM1* among *Triticeae* species. The ratio of non-synonymous to synonymous nucleotide exchanges (Ka/Ks) among *PUB15* and *ARM1* genes of four *Triticeae* species was calculated in a stepwise sliding window of 120 nucleotides. Deviation of the mean Ka/Ks ratios of all six pairwise species comparisons from the null-hypothetical value “1” was tested by t-test. NS, not significantly different from 1 ( p>0.01, 2-tailed). **(a-e)**Species binomial abbreviations: *Aet*, *Aegilops tauschii* (wild wheat); *Hv, Hordeum vulgare* (barley); *Sc, Secale cereale* (rye), *Ta, Triticum aestivum* (wheat).

**Table 1.** Sequence overview of *HvPUB15* and its partial duplicate *HvARM1* in the barley genome.

| Identifier | *HvPUB15* | *HvARM1* |
| --- | --- | --- |
| cDNA clone ID | HO23D08 | HO14H18 |
| HarvEST assembly #35 unigene Nr. | 3072 | 3071 |
| Full-coding sequence cDNA Acc. Nr. | AK361754 | AK371875 |
| Morex WGS contig Acc. Nr.<sup>a</sup> | CAJW010005672 | CAJX010121345 |
| Barke WGS contig Acc. Nr.<sup>a</sup> | CAJV010187631 | CAJV010188692 |
| Bowman WGS contig Acc. Nr.<sup>a</sup> | CAJX010851782 | CAJX010121345 |
| High confidence barley gene ID<sup>a</sup> | HORVU3Hr1G113910 | HORVU3Hr1G081380 |
| Chromosome<sup>b</sup> | 3HL | 3HL |
| Position (Mbp)<sup>b</sup> | 689,57 | 594,73 |
| Syntenic to *B.distachyon*, *O.sativa*,<br>*S.bicolor* | No | No |
| Syntenic to *A. tauschii* | Yes | No |

<sup>a</sup>Most significant BlastN result with 99-100% identity to genomic sequence of barley (http://webblast.ipk-gatersleben.de/barley/).
<sup>b</sup>Based on high-confidence (HC)-gene mapping of the barley reference sequence (Mascher *et al.*, 2017).

### Allelic variants of HvARM1

The phylogenetic and functional (see below) data of *ARM1* suggest that the gene is under selection for maintaining a quantitative level of resistance of *Triticeae* species to powdery mildew infection. We therefore analyzed gene variants (alleles) in a diverse collection of barley genotypes (supporting Tables S1 and S2) for significant association with powdery-mildew interactions. To address this question further we carried out an association-genetic analysis of single nucleotide polymorphism (SNP) and gene-haplotypes with powdery-mildew resistance or-susceptibility. As shown in Table 2this led to the identification of SNP-as well as gene-haplotype polymorphisms in two diverse, world-wide collections of barley landraces and cultivars that were significantly associated with quantitative powdery mildew resistance. No association of *HvPUB15* gene variants with the same trait was found in any population (Table 2). This result strengthens the point that *HvARM1* - despite its partial nature - represents a functional gene protecting barley from powdery mildew attack, whereas the cellular functions of *HvPUB15* are probably more complex.

**Table 2.**
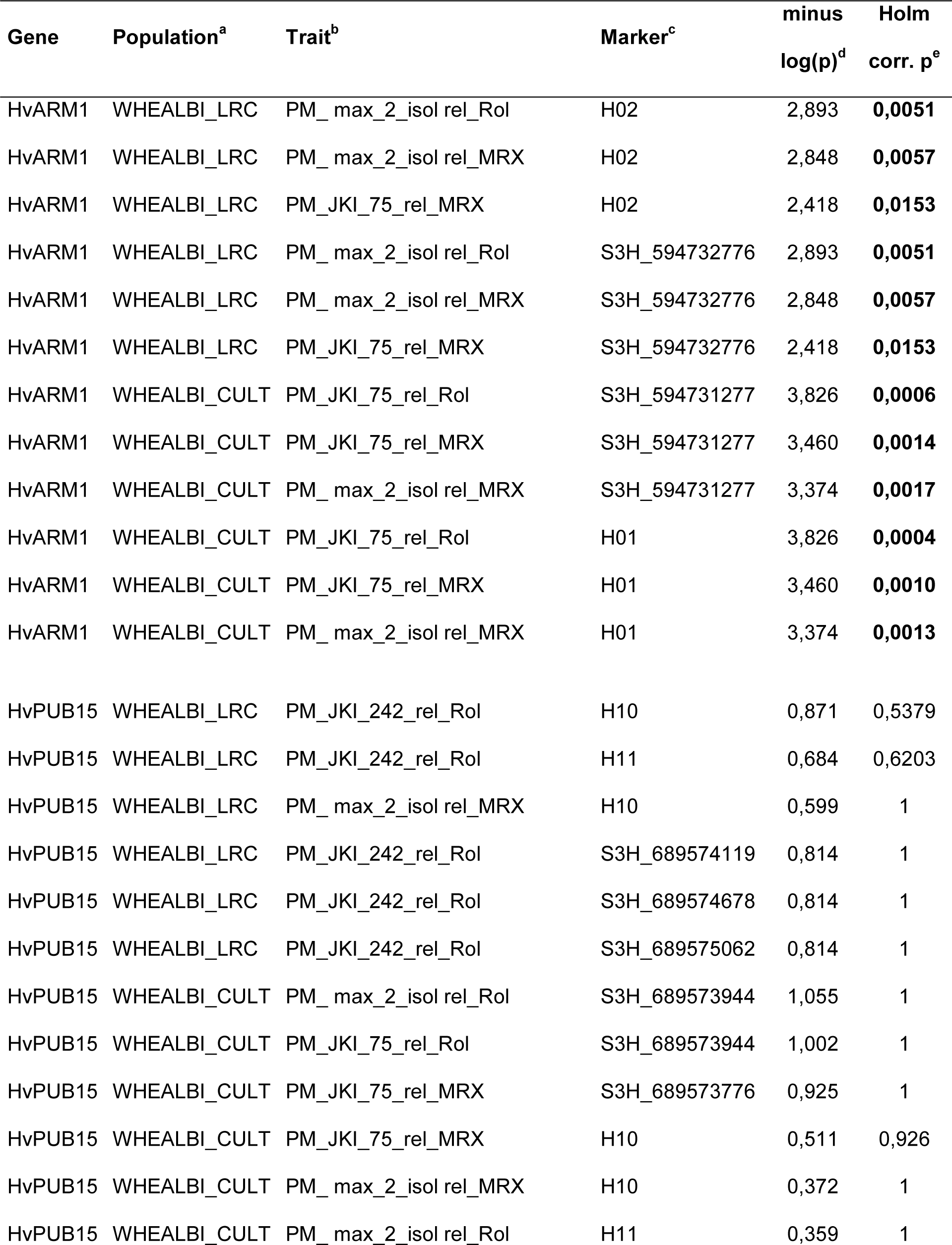
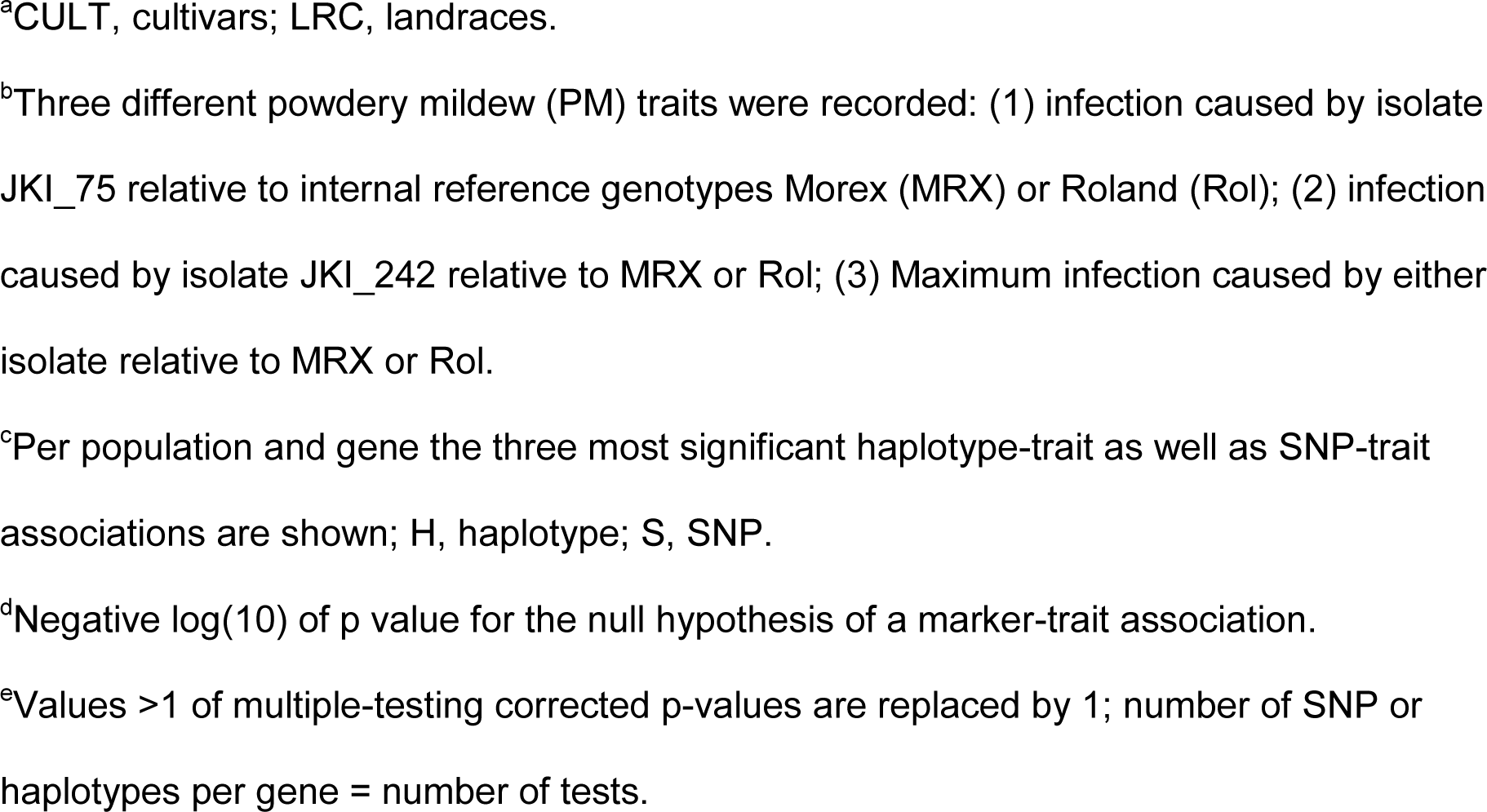
Marker-trait associations of *HvARM1* and *HvPUB15* in diverse collections of cultivated *H. vulgare ssp. vulgare*.

| Gene | Population<sup>a</sup> | Trait<sup>b</sup> | Marker<sup>c</sup> | minus<br>log(p)<sup>d</sup> | Holm<br>corr. p<sup>e</sup> |
| --- | --- | --- | --- | --- | --- |
| HvARM1 | WHEALBI_LRC | PM_max_2_isol rel_Rol | H02 | 2,893 | **0,0051** |
| HvARM1 | WHEALBI_LRC | PM_max_2_isol rel_MRX | H02 | 2,848 | **0,0057** |
| HvARM1 | WHEALBI_LRC | PM_JKI_75_rel_MRX | H02 | 2,418 | **0,0153** |
| HvARM1 | WHEALBI_LRC | PM_max_2_isol rel_Rol | S3H_594732776 | 2,893 | **0,0051** |
| HvARM1 | WHEALBI_LRC | PM_max_2_isol rel_MRX | S3H_594732776 | 2,848 | **0,0057** |
| HvARM1 | WHEALBI_LRC | PM_JKI_75_rel_MRX | S3H_594732776 | 2,418 | **0,0153** |
| HvARM1 | WHEALBI_CULT | PM_JKI_75_rel_Rol | S3H_594731277 | 3,826 | **0,0006** |
| HvARM1 | WHEALBI_CULT | PM_JKI_75_rel_MRX | S3H_594731277 | 3,460 | **0,0014** |
| HvARM1 | WHEALBI_CULT | PM_max_2_isol rel_MRX | S3H_594731277 | 3,374 | **0,0017** |
| HvARM1 | WHEALBI_CULT | PM_JKI_75_rel_Rol | H01 | 3,826 | **0,0004** |
| HvARM1 | WHEALBI_CULT | PM_JKI_75_rel_MRX | H01 | 3,460 | **0,0010** |
| HvARM1 | WHEALBI_CULT | PM_max_2_isol rel_MRX | H01 | 3,374 | **0,0013** |
| HvPUB15 | WHEALBI_LRC | PM_JKI_242_rel_Rol | H10 | 0,871 | 0,5379 |
| HvPUB15 | WHEALBI_LRC | PM_JKI_242_rel_Rol | H11 | 0,684 | 0,6203 |
| HvPUB15 | WHEALBI_LRC | PM_max_2_isol rel_MRX | H10 | 0,599 | 1 |
| HvPUB15 | WHEALBI_LRC | PM_JKI_242_rel_Rol | S3H_689574119 | 0,814 | 1 |
| HvPUB15 | WHEALBI_LRC | PM_JKI_242_rel_Rol | S3H_689574678 | 0,814 | 1 |
| HvPUB15 | WHEALBI_LRC | PM_JKI_242_rel_Rol | S3H_689575062 | 0,814 | 1 |
| HvPUB15 | WHEALBI_CULT | PM_max_2_isol rel_Rol | S3H_689573944 | 1,055 | 1 |
| HvPUB15 | WHEALBI_CULT | PM_JKI_75_rel_Rol | S3H_689573944 | 1,002 | 1 |
| HvPUB15 | WHEALBI_CULT | PM_JKI_75_rel_MRX | S3H_689573776 | 0,925 | 1 |
| HvPUB15 | WHEALBI_CULT | PM_JKI_75_rel_MRX | H10 | 0,511 | 0,926 |
| HvPUB15 | WHEALBI_CULT | PM_max_2_isol rel_MRX | H10 | 0,372 | 1 |
| HvPUB15 | WHEALBI_CULT | PM_max_2_isol rel_Rol | H11 | 0,359 | 1 |

<sup>a</sup>CULT, cultivars; LRC, landraces.
<sup>b</sup>Three different powdery mildew (PM) traits were recorded: (1) infection caused by isolate JKI\_75 relative to internal reference genotypes Morex (MRX) or Roland (Rol); (2) infection caused by isolate JKI\_242 relative to MRX or Rol; (3) Maximum infection caused by either isolate relative to MRX or Rol.
<sup>c</sup>Per population and gene the three most significant haplotype-trait as well as SNP-trait associations are shown; H, haplotype; S, SNP.
<sup>d</sup>Negative log(10) of p value for the null hypothesis of a marker-trait association.
<sup>e</sup>Values >1 of multiple-testing corrected p-values are replaced by 1; number of SNP or haplotypes per gene = number of tests.

### Function of HvARM1 during powdery-mildew attack

To validate the TIGS effect of *HvARM1* in the nonhost interaction with *Bgt* (Douchkov et al., 2014), and to further assess its role in the interaction with the adapted *Bgh* we generated transgenic plants with silenced Hv*ARM1*. Approximately 25% of progeny from T0 primary transformants died after a few weeks suggesting lethality caused by the homozygous transgene locus, in line with the failure to identify homozygous T2 or T3 lines(Figure 3c, pot Nr. 1 in left-hand panel). It is known that homozygous transgenic plants usually exhibit strongest transgene effects, and this may have caused off-target silencing ofthe related, housekeeping *HvPUB15* gene as suggested from *in silico* off-target prediction ofthe RNAi transgene by the si-Fi software. Using default settings including end-stability difference and target-site accessibility thresholds, the software predicted 33 and 6 efficient21nt siRNAs for *HvARM1* and *HvPUB15*, respectively (supporting Figure S3). The suspected toxicity of *HvPUB15* off-target silencing in homozygous RNAi plants is supported by observations in *Oryza sativa* (rice), where a detrimental effect of knock-out mutation of Os*PUB15* was described including seedling lethality and severe growth retardation (Park et al., 2011). On the other hand, the normally developing T3 transgenic lines consistently exhibited silencing of *HvARM1*whereas no evidence for a reduction of *HvPUB15*mRNA levels was found (Figure 3a).

**Figure 3:**
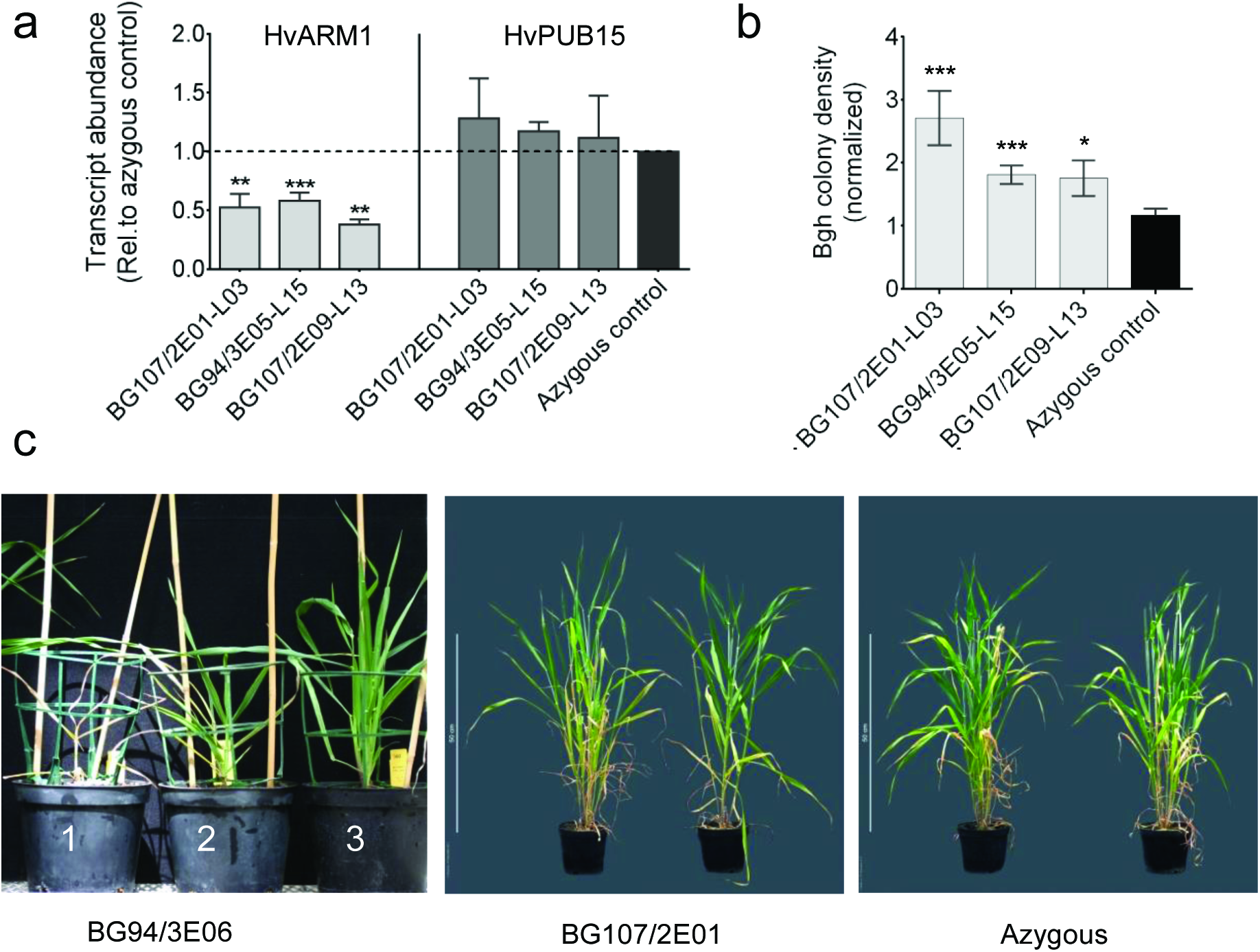
Silencing of *HvARM1* and *HvPUB15* in *H*. vulgare affects quantitative resistance against *B. graminis f. sp. hordei*. **(a)**Transcript abundance of the *HvARM1* target gene and its possible off-target *HvPUB15* was determined by RT-qPCR in RNA from leaves of non-inoculated plants. Normalized transcript abundance relative to the *HvUBC* reference gene encoding an E2 ubiquitin conjugating enzyme (see “Materials and Methods”) was further normalized to the mean value of azygous segregants (set to “1”). Single T2-plant-derived T3 sister lines are indicated by small letters a or b. Mean values ± SE from 3 biological replicates (batches of plants sown on different dates) are shown. **(b)**Detached second leaves of T3 transgenic barley RNAi plants were inoculated with *Bgh* and infection was assessed microscopically 48 hours after inoculation. Data represent normalized colony density (number/cm^2^/median of azygous control per experiment) ± SE from 2-3 biological replications. Differences between transgenic events and azygous plants are indicated by asterisks. *, p<0.05; ***, p<0.0005 (student’s t-test; two-tailed). **(c)**Growth phenotypes of *HvARM1*-silenced transgenic seedlings from event BG94/3E06 exhibiting more severe seedling lethality (not used for *Bgh*-interaction phenotyping in T3 generation) and of adult plants from event BG107/2E01 that was used for *Bgh*-resistance tests. Plant numbers 1, 2 and 3 show examples of seedlings with lethal, growth-retarded and wildtype-like growth phenotypes.

Because *HvARM1* was discovered in a TIGS screen for attenuated NHR we first tested T3 progeny of three selected events for susceptibility to *Bgt* (Table 3). Although there was a considerable variability between individuals per line, two lines exhibited approximately five-fold higher susceptibility to the non-adapted fungus as compared to the control group of azygous segregant plants. In general, azygous plants are considered as better controls since they had undergone the same transformation procedures and lost the transgenic construct by segregation. As seen in Table 3, the transformation procedure apparent had an impact on the *Bgt*interaction because the azygous control group was on average more susceptible than Golden Promise wildtype plants. Figure 3b shows that the three selected events were also more susceptible to *Bgh*, compared to a population of control plants consisting of azygous segregants plus progeny from three azygous individuals identified in the T2 generation.

**Table 3.** Reduced nonhost resistance of *HvARM1*-silenced transgenic *H*. vulgare plants against *B. graminis f.sp. tritici*.

| Genotype<sup>a</sup> | Colonies/leaf segment<sup>b</sup> | N<sup>c</sup> | p-value<sup>d</sup> |
| --- | --- | --- | --- |
| Wildtype (G. Promise) | 0.380 ± 0.307 | 25 | <0.0001 |
| Azygous control | 3.332 ± 0.686 | 78 |  |
| BG107/2E01-Line-3 | 4.153 ± 1.010 | 37 | 0.2518 |
| BG94/3E05-Line-15&20<sup>e</sup> | 17.510 ± 7.582 | 47 | 0.0345 |
| BG107/2E09-Line-13 | 16.310 ± 7.067 | 28 | 0.0392 |

<sup>a</sup>Transgenic- and null-segregant plants are from the T3 generation.
<sup>b</sup>Microcolonies were counted 7 days after inoculation under the light microscope. Total number of colonies per leaf segments that were cut to the same length. Mean values ± SEM.
<sup>c</sup>Number of leaf segments (plants).
<sup>d</sup>t-test (1-tailed) against the azygous control, asking for enhanced colony number.
<sup>e</sup>Data from T3 progeny of two T2 sister lines were pooled in order to reach a sufficiently large number of tested individuals.

Bombardment with strictly gene-specific RNAi-and with over-expression (OEX) constructs for a direct comparison of altered *HvARM1 versus HvPUB15* expression levels revealed that gene-specific *HvARM1* silencing increased the relative susceptibility index (SI) to *Bgh*, in line with the super-susceptibility observed in stable transgenic barley T3 plants (Table 4). On the other hand, we found no significant effects of altering *HvPUB15* mRNA levels on the interaction of transformed cells with *Bgh* again indicating more complex, homoeostatic rather than defense-related functions of the encoded protein. Following powdery mildew inoculation, endogenous transcript levels of *HvARM1* in peeled leaf epidermis were more strongly up-regulated above a basal level of expression compared to *HvPUB15* (supporting Figure S4), which also suggests a defense-related role of *HvARM1*.

**Table 4.**
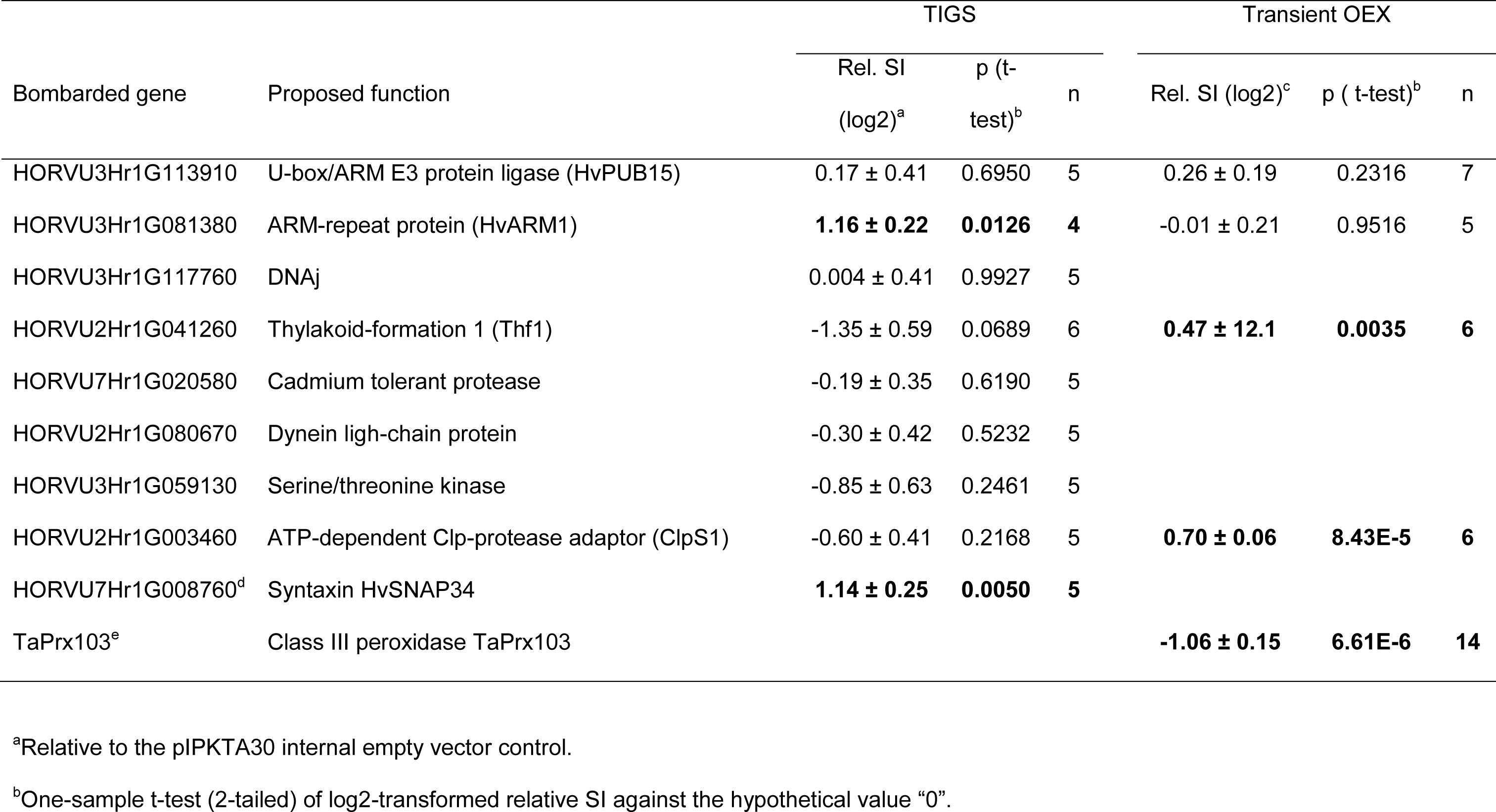
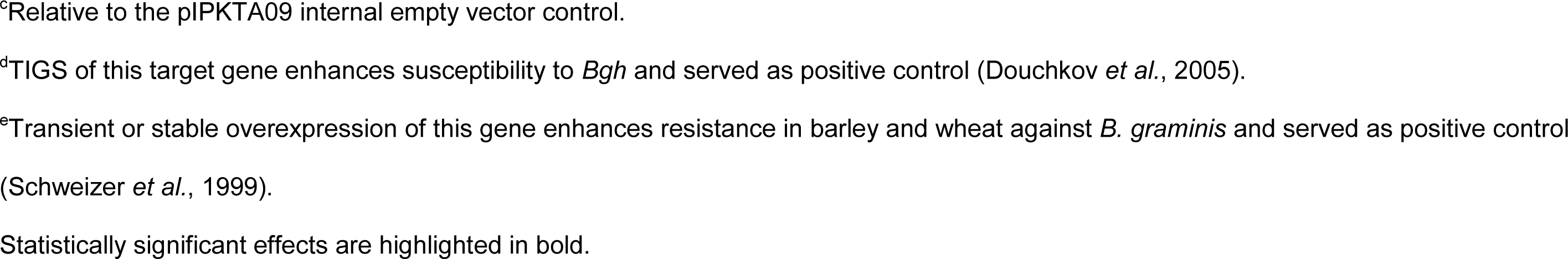
Effect of TIGS and transient over-expression of *HvARM1* and genes encoding its interacting proteins on QR against *B. graminis f.sp. hordei*.

| Bombarded gene | Proposed function | TIGS |  |  | Transient OEX |  |  |
| --- | --- | --- | --- | --- | --- | --- | --- |
|  |  | Rel. SI<br>(log2)<sup>a</sup> | p (t-test)<sup>b</sup> | n | Rel. SI (log2)<sup>c</sup> | p (t-test)<sup>b</sup> | n |
| HORVU3Hr1G113910 | U-box/ARM E3 protein ligase (HvPUB15) | 0.17 ± 0.41 | 0.6950 | 5 | 0.26 ± 0.19 | 0.2316 | 7 |
| HORVU3Hr1G081380 | ARM-repeat protein (HvARM1) | **1.16 ± 0.22** | **0.0126** | **4** | -0.01 ± 0.21 | 0.9516 | 5 |
| HORVU3Hr1G117760 | DNAj | 0.004 ± 0.41 | 0.9927 | 5 |  |  |  |
| HORVU2Hr1G041260 | Thylakoid-formation 1 (Thf1) | -1.35 ± 0.59 | 0.0689 | 6 | **0.47 ± 12.1** | **0.0035** | **6** |
| HORVU7Hr1G020580 | Cadmium tolerant protease | -0.19 ± 0.35 | 0.6190 | 5 |  |  |  |
| HORVU2Hr1G080670 | Dynein ligh-chain protein | -0.30 ± 0.42 | 0.5232 | 5 |  |  |  |
| HORVU3Hr1G059130 | Serine/threonine kinase | -0.85 ± 0.63 | 0.2461 | 5 |  |  |  |
| HORVU2Hr1G003460 | ATP-dependent Clp-protease adaptor (ClpS1) | -0.60 ± 0.41 | 0.2168 | 5 | **0.70 ± 0.06** | **8.43E-5** | **6** |
| HORVU7Hr1G008760<sup>d</sup> | Syntaxin HvSNAP34 | **1.14 ± 0.25** | **0.0050** | **5** |  |  |  |
| TaPrx103<sup>e</sup> | Class III peroxidase TaPrx103 |  |  |  | **-1.06 ± 0.15** | **6.61E-6** | **14** |

<sup>a</sup>Relative to the pIPKTA30 internal empty vector control.
<sup>b</sup>One-sample t-test (2-tailed) of log2-transformed relative SI against the hypothetical value “0”.
<sup>c</sup>Relative to the pIPKTA09 internal empty vector control.
<sup>d</sup>TIGS of this target gene enhances susceptibility to *Bgh* and served as positive control (Douchkov *et al.*, 2005).
<sup>e</sup>Transient or stable overexpression of this gene enhances resistance in barley and wheat against *B. graminis* and served as positive control (Schweizer *et al.*, 1999).
Statistically significant effects are highlighted in bold.

### Localization and protein interactions of HvPUB15 and HvARM1

Fusion proteins of HvARM1 and HvPUB15 with the yellow fluorescent protein (YFP) showed a similar fluorescence pattern as non-fused YFP suggesting nucleo-cytoplasmic localization (supporting Figure S5, panels a-c), in agreement with the localization of the proposed rice orthologue OsPUB15 (Park *et al.*, 2011). Because the presence of the conserved ARM protein-protein interaction domain in HvARM1 suggests binding to other barley protein(s), we carried out a yeast-two-hybrid screening in a prey library from *Bgh*-attacked barley leaves using HvARM1 as bait. This led to the identification of six barley proteins that interacted strongly and reproducibly with HvARM1 (Figure 4a and supporting Table S3). Out of these six candidates the homologs of the Clp-protease adaptor protein ClpS1 and the thylakoid formation 1 protein THF1 of *A. thaliana* (Huang *et al.*, 2006, Nishimura *et al.*, 2013) were also strongly interacting with HvPUB15 (Figure 4b), which suggests them as ubiquitination substrates of the E3 ligase. By using an *in vitro* ubiquitination assay we could show that HvPUB15 has ubiquitin ligase activity (supporting Figure S6). Because HvPUB15 catalyzed the polymerization of ubiquitin chains rather than auto-ubiquitination it might be an E4-rather than an E3-ligase (Koegl *et al.*, 1999). The possible involvement of all six HvARM1-interacting proteins in mediating QR or susceptibility to *Bgh* was tested by TIGS, which revealed a trend for reduced susceptibility by silencing of *HvThf1* (Table 4). Transient OEX of *HvThf1*resulted in significantly enhanced susceptibility to *Bgh* thereby proposing the encoded potential HvPUB15 substrate protein to function as host susceptibility factor. Transient OEX of the second proposed HvPUB15 substrate, *HvClpS1*, also enhanced susceptibility to *Bgh* thereby indicating a role as host susceptibility factor, too, although there was no significant effect of *HvClpS1* silencing. Co-localization experiments of HvThf1-YFP and HvClpS1-YFP C-terminal fusion proteins with the plastid marker Rubisco small subunit (Nelson *et al.*, 2007) confirmed their expected plastid localization (supporting Figure S5, panels d-k).

**Figure 4:**
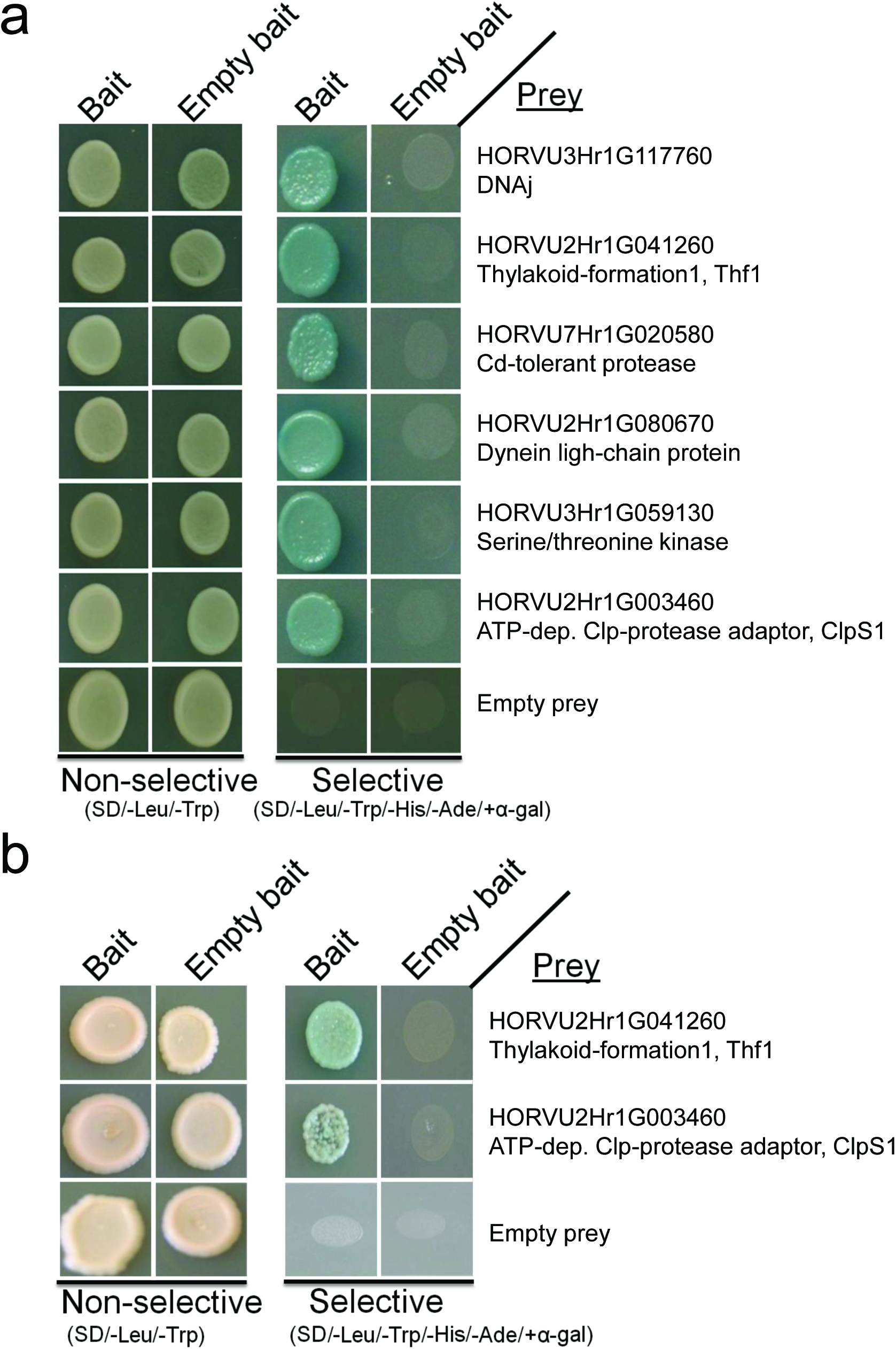
Proteins of *H. vulgare* interacting with HvARM1 and HvPUB15 in the yeast-two-hybrid system (Y2H). **(a)**Full-length HvARM1 was used as bait in a Y2H screening of a cDNA library derived from *Bgh*-attacked barley leaves. Growth of yeast on SD-Leu/-Trp confirms the presence of both bait and prey vectors for protein expression. Growth on SD-Leu/-Trp/-His/-Ade indicates protein-protein interaction. No growth of empty bait vector + candidate prey confirms the absence of autoactivation of any prey construct for six final candidates. **(b)**Two of the six candidate protein interactors of HvARM1 also interact with full-length HvPUB15.

*In vivo* interaction of HvARM1, HvPUB15 and HvPUB15^ARM^(the ARM-domain of HvPUB15) with HvThf1 and HvClpS1 was assessed by split-YPF bimolecular functional complementation (BiFC) assays in *Agrobacterium*-infiltrated *Nicotiana bentaminiana* leaves and by co-immunoprecipitation in *A. thaliana* protoplasts. Figure 5 shows that the transient co-expression of HvPUB15 or HvPUB15^ARM^with either HvClpS1 or HvThf1 gave rise to BiFC (YFP) signals primarily in epidermal cells. The localization patterns of the fluorescence signals indicated that the proteins interacted in the cytoplasm, which was confirmed by the absence of co-localization with the plastid marker protein 35S:SSU_1-79_-mCherry (Rajaraman, unpublished result). The BiFC signals were abolished or strongly reduced by using the U-box mutant HvPUB15^P245A^as interaction partner, suggesting specificity of the interaction. HvARM1 also interacted with HvClpS1 and HvThf1. Moreover, interactions were observed between HvPUB15 and HvARM1, and here, the HvPUB15 U-box mutation increased BiFC signals instead of reducing them. For additional controls and quantitative fluorescence data see supporting Figures S7 and S8. Co-immunoprecipitation experiments in *A. thaliana* protoplasts of cMyc-tagged HvARM1 and HvPUB15^ARM^together with either HvThf1 or HvClpS1 confirmed *in vivo* interaction of HvPUB15^ARM^with HvThf1, HvARM1 with HvThf1, and HvARM1 with HvClpS1 (Figure 6).

**Figure 5:**
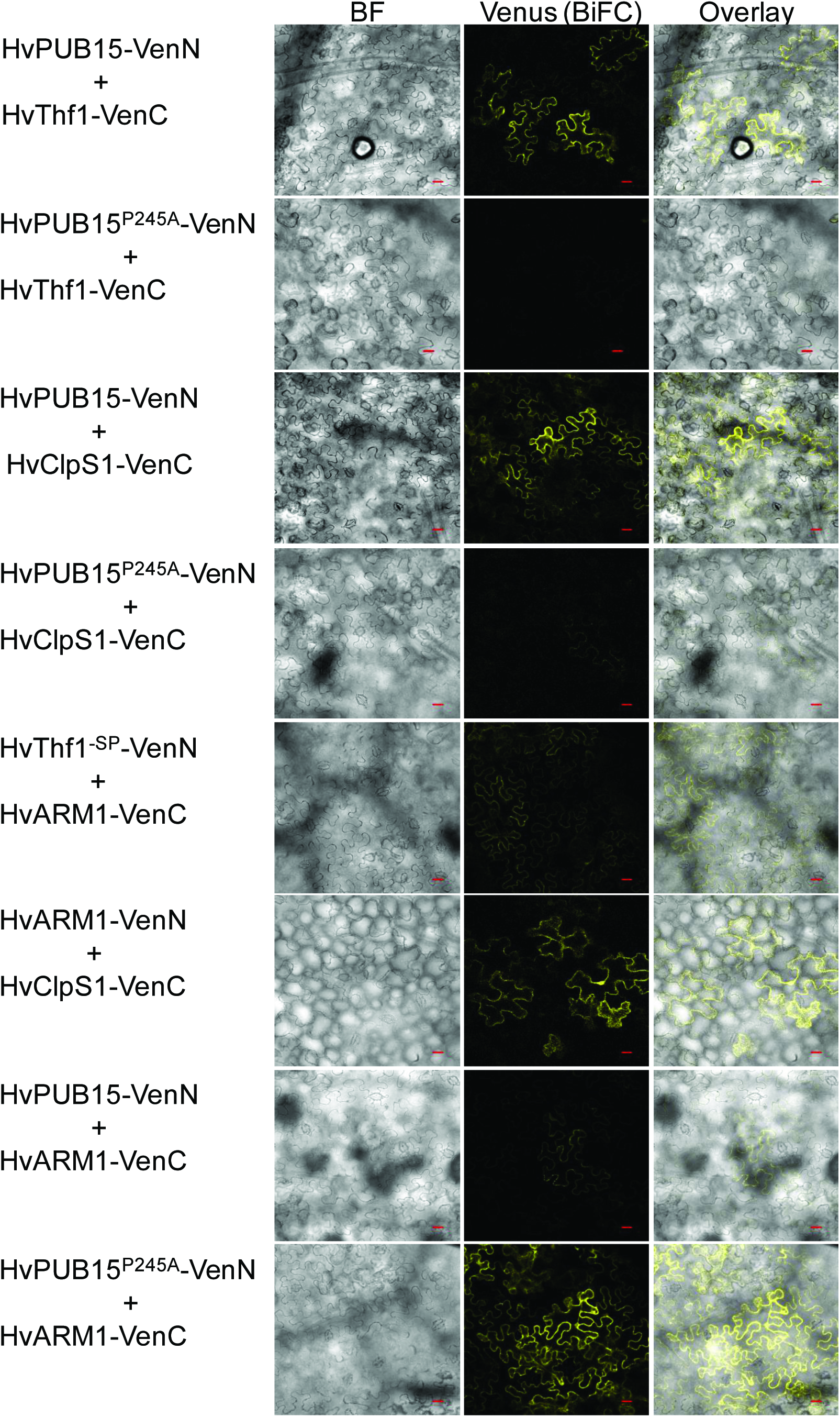
Bimolecular functional complementation (BiFC) of YFP by *H. vulgare* proteins interacting with HvARM1 and HvPUB15 *in vivo*. BiFC by HvPUB15- and HvARM1-interacting proteins in *N. benatminiana* leaves after infiltration of *A. tumefaciens* strains carrying the protein interaction partners fused C-terminally to split halves of YFP. BF, bright field;-SP, with deleted N-terminal plastid import signal; VenN, N-terminal half of the stabilized YFP version “Venus”; VenC, C-terminal half of the YFP “Venus”. Scale bars, 20 µm.

**Figure 6:**
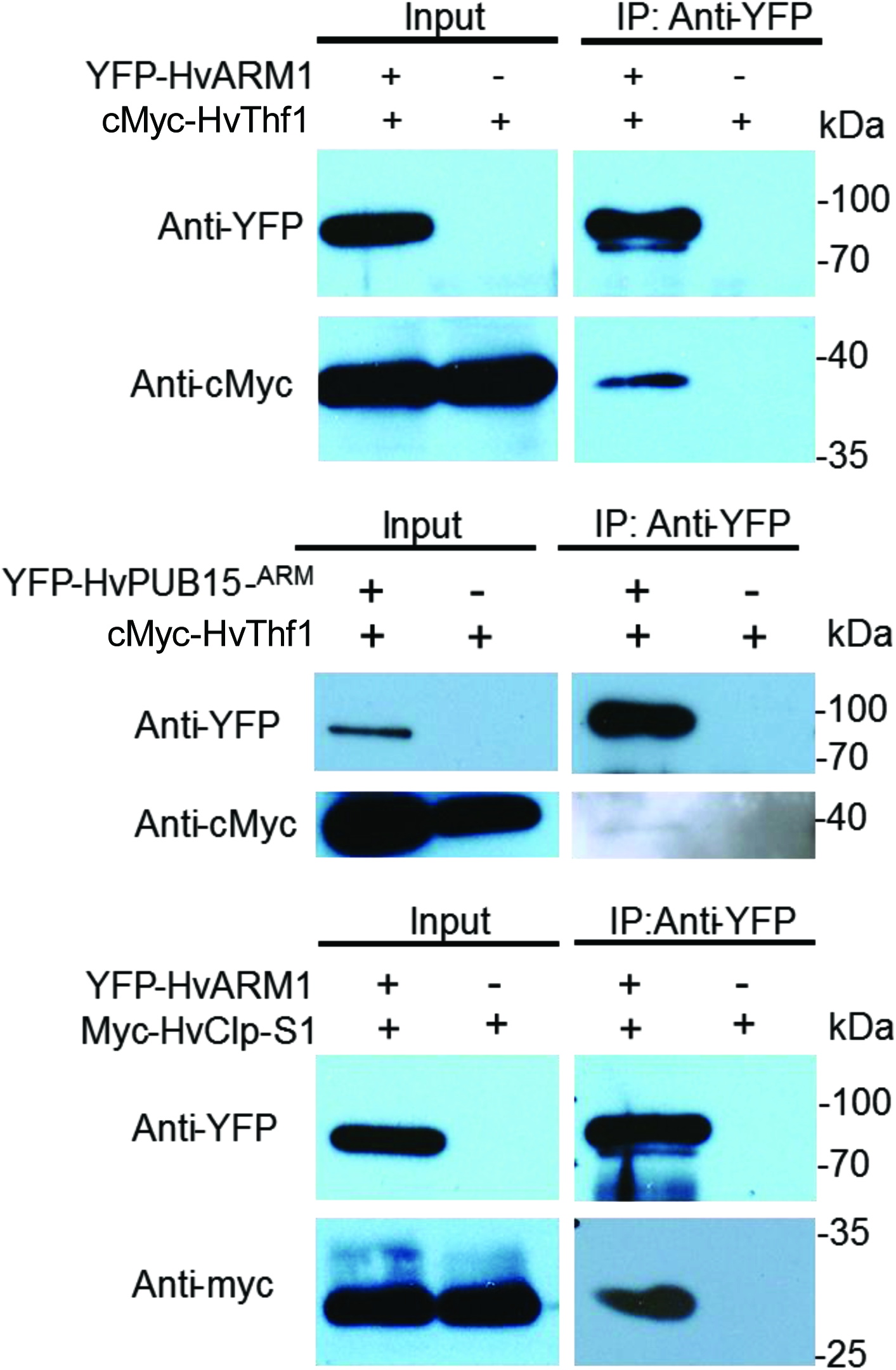
Co-immunoprecipitation (Co-IP) of *H. vulgare* proteins interacting with HvARM1 and HvPUB15 *in vivo*. Co-IP of antibody-tagged barley proteins in *A. thaliana* mesophyll protoplasts. YFP-fused *HvARM1* and *HvPUB15* were co-expressed with cMyc-tagged HvThf1 and *HvClpS1*, respectively for each interaction. Co-IP was performed using anti-YFP antibodies and total proteins extracted from *A. thaliana* protoplasts.

The two HvPUB15 interacting proteins HvThf1 and HvClpS1 may be *in vivo*ubiquitination substrates. This possibility was tested by transient over-expression of *HvPUB15* together with either HvThf1:YFP or HvClpS1:YFP, followed by the quantification of YFP-fluorescing cells 24 hours after the bombardment. As shown in Figure 7, co-expression with *HvPUB15* significantly reduced the number of HvThf1:YFP-but not HvClpS1:YFP-fluorescing cells suggesting that HvThf1 is an *in vivo* substrate to HvPUB15. Taken together, the data suggest *in vivo* interaction of HvPUB15 with the putative substrate proteins HvClpS1 and HvThf1, whereby the presence of an intact U-box stabilized the interaction. In addition, HvARM1 as well as the ARM-domain of HvPUB15 interacted with HvClpS1 and HvThf1, too. One of the HvPUB15-interacting proteins, HvThf1, was identified as potential ubiquitination substrate and as host susceptibility factor to *Bgh*.

Does the partially duplicated gene pair of PUB15 and ARM1 represent a unique event in *Triticeae* genome evolution, or could we find indications for additional partial gene duplicates with putative functions? To address this question we conducted a genome-wide search for full-length cDNA sequences with high sequence similarity but clearly different length of their longest open reading frames. Starting with a library of 23,614 full-length cDNA sequences (http://barleyflc.dna.affrc.go.jp/bexdb/), we found 1154 matching cDNA pairs with a sequence identity of 80-99% and an alignment-to-shorter-gene length ratio of at least 0.8, thereby excluding pairs of non-partial genes just sharing functional domains. A subsequent tBlastx analysis of these pairs revealed 205 pairs with a length difference of matching open reading frames of >25%. After further filtering steps to exclude non-spliced transcripts, chimeric-as well as partial clones, we identified 11 expressed pairs of putative, partially duplicated genes including *HvPUB15/HvARM1* (supporting Table S4). A majority of these are localized at non-tandem positions (5 or more gene models apart from each other, or on different chromosomes). Interestingly, although this was not a criterion for their selection, transcripts from the partially duplicated genes appeared to be more frequently regulated by powdery mildew attack (Figure 8a). This suggests that partial gene duplicates might be preferentially selected for new functions in stress responses such as pathogen attack. As another example of a partially duplicated gene and protein pair besides HvPUB15/HvARM1, we show the alignment of a DUF 4228 protein of unknown function (Figure 8b and c). This pair, which is located 5 gene models apart from each other on chromosome 7H, is characterized by a clear gain in transcript regulation of the shorter version and by a perfect triple repeat of a highly hydrophilic motif in the duplicated part of the protein. Its functional analysis in pathogen-attacked barley will be a future challenge.

**Figure 8:**
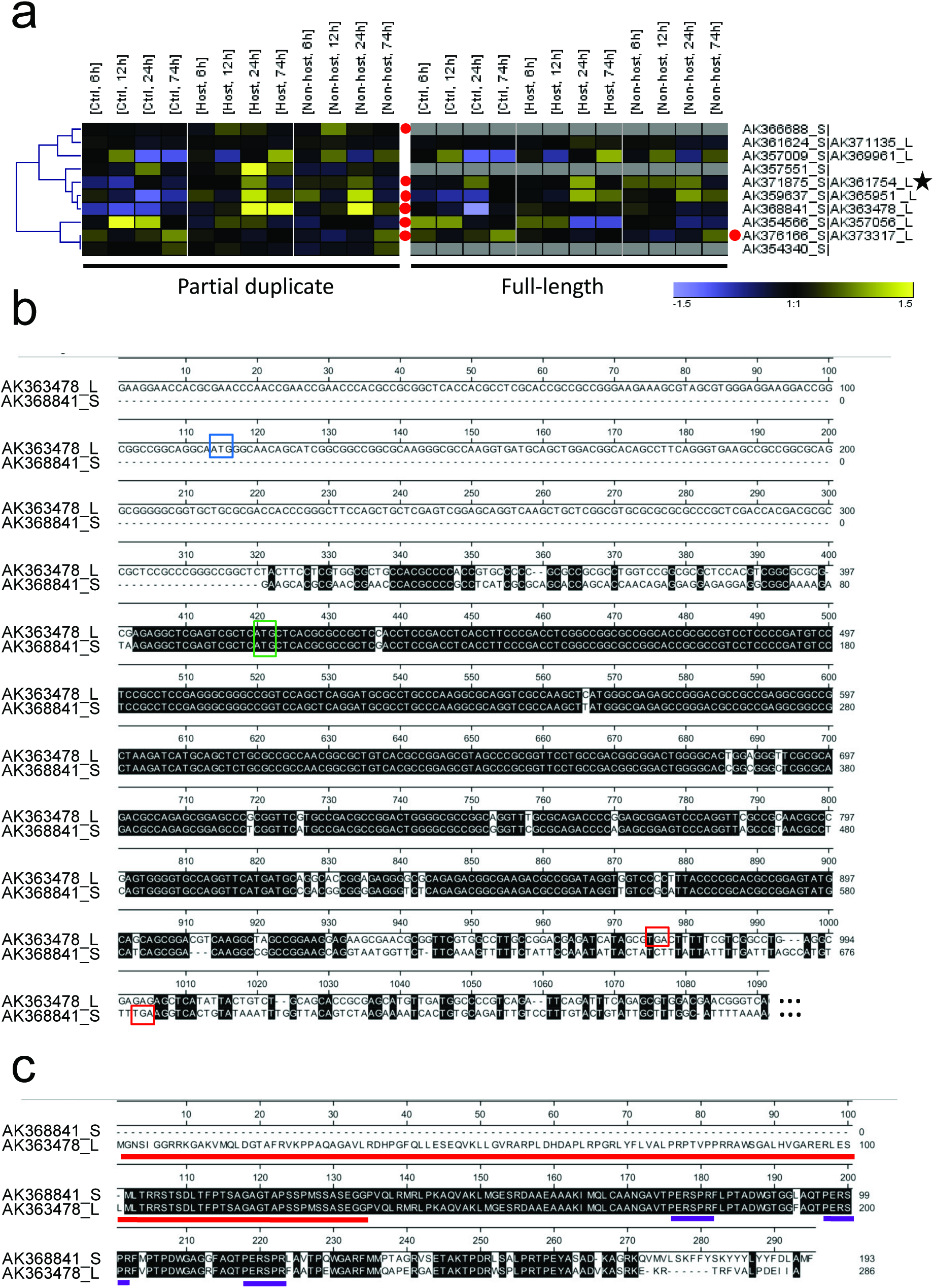
Genome-wide search for expressed, partial gene duplicates. **(a)**Transcript regulation in peeled barley leaf epidermis by Bgh (host) or Bgt (nonhost). Total RNA was isolated at the time indicated after inoculation and subjected to hybridization to the Barley Gene Expression Array of Agilent. For the link of full-length cDNA accession No. to Agilent probe IDs, see supporting Table S2. Transcript data have been submitted to ArrayExpress (Acc. E-MTAB-2916). Hierarchical clustering of gene-median-centered, normalized signal intensities is shown. The color scale ranges from log(2)-1.5 to 1.5. Mean signal intensities from three independent inoculation experiments are shown. **(b)**cDNA Alignment example of a DUF4228 protein and its proposed partial duplicate. **(c)**Protein alignment of the translated sequences of cDNAs shown in (b). Red and violet bars indicate the DUF4228 domain and a perfect repeat of a highly hydrophilic protein motif, respectively.

## DISCUSSION

The genomes of four species of the *Triticeae* tribe of grasses contain *ARM1*, a partial copy of a U-box/ARM-repeat E3 ligase closely related to the rice protein OsPUB15 (Park *et al.*, 2011). The rice genome also contains a number of “ARM-repeat only” genes but none of them appears to represent a partial copy of Os*PUB15*, because BlastN analysis of the ARM-repeat region of Os*PUB15* (pos. 2000-2952 in cDNA Acc. AK10655) at NCBI did not produce significant hits for any other rice gene. By contrast, the same query sequence revealed Hv*ARM1* as the most significant hit (86% identity) in barley. Sequence analysis in cultivated barley and rye, and in two diploid wild wheat species suggest a monophyletic origin of *ARM1.* In a TIGS screening for genes required for NHR in barley, we discovered *Rnr5* encoding HvARM1 as potential factor for resistance to the non-adapted powdery mildew fungus *Bgt.* Here we show that Hv*ARM1* is not only involved in NHR but is also a factor for QR to adapted powdery mildew fungi in barley and wheat.

The large and highly repetitive genomes of the cultivated *Triticeae* species barley, wheat and rye appear to be particularly rich in gene-like sequences including partial duplicates, and most of them were classified as putative pseudogenes (Akhunov *et al.*, 2013, Wicker *et al.*, 2011, Bauer *et al.*, 2017). The classification criteria for these putative pseudogenes were (i) non-syntenic map positions among grasses and (ii) unique occurrence in one species or in one of the three sub-genomes of hexaploid wheat. Illegitimate meiotic crossing over and subsequent sequence capture by transposable elements, as well as random-sequence insertion during non-homologous end joining for double-strand break DNA repair are the two proposed major events leading to non-tandem (partial) gene duplicates (Katju & Lynch, 2003). By contrast, sub- or neo-functionalized, expressed and full-length gene duplicates often exist as tandemly repeated gene pairs or clusters of genes, as a result of unequal crossover during meiosis that often followed by gene-conversions (Himmelbach *et al.*, 2010, Leister, 2004). As shown in Table 1, the full-length genes Hv*PUB15* and Aet*PUB15* share syntenic map positions on the long arm of homologous chromosome group 3 (Luo *et al.*, 2013, Mascher *et al.*, 2013). The partially duplicated Hv*ARM1* gene was mapped at a distance of approximately 95 Mbp from Hv*PUB15* on chromosome 3H, and all four analyzed *Triticeae ARM1* genes contain a non-repetitive, unknown sequence in exon 1 that is not present in the corresponding *PUB15*-like genes. Taken together this suggests that an event of DNA double-strand break repair in a common ancestor of the four *Triticeae* species gave rise to *ARM1*.

However, the results presented here suggest that *ARM1* escaped pseudogenization and took over a new biological function in defense against powdery mildew fungi and perhaps other pathogens: First, the genomes of four species belonging to three different *genera* maintained the partial gene copy with a high degree of sequence conservation at the ARM-repeat region. Second, in all four species *ARM1* is supported by perfectly matching EST sequences or other transcriptome data, demonstrating that the corresponding genes are actively transcribed. In barley, transcript-regulation data suggest a gain of function of *HvARM1* in terms of a more pronounced pathogen-induced accumulation in the epidermis compared to *HvPUB15* (supporting Figure S4). Third, all *ARM1* sequences are characterized by intact open reading frames starting approximately in the middle part of the PUB15 protein and extending to its C-terminus. Forth, allelic variants of *HvARM1* were found to be significantly associated with the severity of powdery-mildew infection in collections of locally adapted barley landraces and diverse cultivars (Table 2 and supporting Table S1). In both collections the most significant SNP were associated with clear and statistically significant differences in Bgh infection (41% versus 57%, p=0.00037 in WHEALBI_LRC; 35% versus 51%, p=0.023 in WHEALBI_CULT). The significant SNP in the landrace collection was located in the 5’ non-translated region of the HvARM1 transcript whereas the significant SNP in the cultivar collection causes a Glycine to Valine change at position 437 of the encoded protein. The cultivars carrying the corresponding significant, resistance-associated haplotype H01 were derived from very different regions of the world and therefore, probably not similar by descent. The complete absence of association of HvPUB15 alleles with *Bgh* infection furthermore supports the view that the E3 ligase primarily has important housekeeping functions such as quality control and turning over of plastid-localized proteins (Woodson *et al.*, 2015), with no adaptation flexibility during pathogen co-evolution.

Functional tests of *HvARM1* by gene silencing in barley and by transient over-expression in wheat suggested a resistance-related role during the interaction with adapted and non-adapted powdery mildew fungi. Similar to the results presented here, a resistance-enhancing effect was found by over-expression of the ARM domain of the *AtPUB13* gene in *A. thaliana*, which is involved in protein degradation of the flagellin PAMP receptor FLS2, (Zhou *et al.*, 2015). Transgenic plants phenocopied the *atpub12/13* double-mutant effect of enhanced pathogen resistance. The results presented here for *ARM1* genes of *Triticeae*species suggest that plants use ARM-domain expression as natural mechanism for enhancing disease resistance. Results in rice also proposed a defense-related role of *OsPUB15* because over-expression caused spontaneous defense responses and increased pathogen resistance (Wang *et al.*, 2015). However, we could not confirm such an activity by TIGS or transient OEX of the barley homologue *HvPUB15* (Table 4). Together with the reported lethality of *OsPUB15* mutations in rice and with the indications of lethal *HvPUB15* off-targeting in homozygous RNAi lines of barley that carry a hairpin construct against *HvARM1*, these results propose *HvPUB15* as a housekeeping gene that is not directly involved in pathogen defense, at least not against *Bgh*.

The HvARM1 and HvPUB15 proteins interacted in yeast and in plants with the plastid-localized proteins HvClpS1 and HvThf1, and both appear to be susceptibility-related factors based on TIGS-and transient over-expression results. Interestingly, transcripts of both *HvClpS1* and *HvThf1* were found to be down-regulated in peeled epidermis by powdery mildew attack, a response that may be expected for susceptibility-related factors (supporting Figure S9 and Table S3). Currently, the evidence for HvThf1 to be relevant for the powdery-mildew interaction is stronger compared to HvClpS1 because only TIGS of *HvThf1* resulted in a (one-sided significant) trend for enhanced resistance, thereby complementing the transient OEX data, and because OEX of HvPUB15 resulted only in a reduction of tagged HvThf1 protein - but not of HvClpS1 protein - amounts. Therefore, we will concentrate the discussion on HvThf1 as interaction partner here. The THF1 protein of *A. thaliana* was not only found to be localized in the plastid stroma but also at its outer membrane facing the cytoplasm where it was proposed to play a role in sugar sensing (Huang *et al.*, 2006). The plastid-internal pool was implicated in degradation of chlorophyll-protein complexes, especially the core complex proteins D1 and D2, during leaf senescence of *A. thaliana* and *Oryza sativa* (Huang *et al.*, 2006, Wang *et al.*, 2004, Yamatani *et al.*, 2013). In both rice and *A. thaliana*, *thf1* mutants exhibited a stay-green phenotype after the onset of dark-induced senescence and were also impaired in adaption to high-light conditions, which resulted in photobleaching probably due to excess electron flux from PSII. In the powdery mildew-affected epidermis containing mainly photosynthetically inactive leucoplasts, sugar sensing for controlled carbohydrate delivery to established *Bgh* haustoria might be more relevant than control of photo-oxidative damage. It is known that powdery mildew-infected cells have a very high demand of energy equivalents and transport large amounts of glucose into haustoria, a process that appears to depend on SWEET sugar transporters and other factors (Chen *et al.*, 2010, Chen *et al.*, 2012, Scholes *et al.*, 1994). Support for the involvement of *Thf1* in disease responses comes from the finding that the closest wheat homolog to *HvThf1*, designated as *TaToxABP1*, acts as binding protein and target of Toxin A that is produced by the necrotrophic, tan-spot fungal pathogen *Pyrenophora tritici-repentis* (Manning *et al.*, 2007). Toxin A-treatment also triggered an oxidative burst in leaves of wheat and barley (Manning *et al.*, 2010, Manning *et al.*, 2007, Pandelova *et al.*, 2012) thereby providing a link of *Thf1* function with ROS control, at least in chloroplasts, and propose a mode of action of Toxin A. Additional support for a relevant role of *Thf1* in plant susceptibilty is provided by results showing interaction of the Thf1 protein with the I2-like coiled-coil (CC) domains of several NB LRR-type resistance proteins (Hamel *et al.*, 2016). One of the well-examined members of this group is the N protein mediating resistance in *Nicotiana tabaccum* against most Tobamoviruses. The authors found that THF1 strongly suppressed N-mediated HR and that the I2-domains of corresponding activated R-proteins interacted with Thf1 in the cytoplasm and thereby, destabilized the protein for degradation. Because cell-death suppression is a hallmark of susceptible interactions with biotrophic pathogens, the proposed function of HvThf1 as susceptibility factor to *Bgh* is in line with the proposed function in *Solanaceaous* plants. The link of proteasomal protein degradation with chloroplast biology has recently been established by reports on the roles of the closest *HvPUB15* homolog in *A. thaliana* designated as *AtPUB4*, and of *AtCHIP*, in plastid quality control and degradation of the caseinolytic plastid peptidase AtClpP4, respectively (Wei et al., 2015; Woodshon et al., 2015). Mutants of *AtPUB4* showed reduced resilience against abiotic stress, indicative of compromised plastid-based control of ROS generation. Plants silenced in- or over-expressing *AtCHIP* exhibited a chlorotic phenotype indicating a strict requirement of accurate control of AtClpP4 levels for cellular homoestasis.

As a model for ARM1 function we propose that the partial duplicate *ARM1* of the ancestral *PUB15* gene was selected as an antagonist of PUB15 thereby disturbing the accurate quality and/or import control of the biotrophic susceptibility factor Thf1 (Figure 9). The antagonistic activity of ARM1 could take place by binding to free Thf1 pre-protein or by interacting with a PUB15/Thf1 proteosomal complex. According to the model, Thf1 inhibition will lead to disturbed sugar sensing and/or plastid functionality, which have to be postulated as requirements to support obligate biotrophic pathogens such as powdery mildew fungi. The possibility of Thf1 as inhibitor of resistance components such as R-proteins (Hamel *et al.*, 2016) is also included in the model and would be compatible with the observed, rapid *HvARM1* TIGS effect on haustorium formation. The role of Thf1 in necrotrophic interactions may be opposite, i.e. resistance-related in terms of preventing pathogen-triggered cell death, which is in line with the observed targeting by the fungal toxin Prt ToxA (Manning *et al.*, 2010). As known for other fungal pathogens, *Bgh* possesses a large arsenal of candidate secreted effector proteins, several of which have been found to contribute to host invasion (Pliego *et al.*, 2013) (Whigham *et al.*, 2015). Therefore it appears also possible that ARM1 acts as a mimic to protect the PUB15 protein from putative *Bgh* effector manipulation (Le Roux *et al.*, 2015, Sarris *et al.*, 2015). However, this possibility is currently not supported by data on ARM1-effector interactions.

**Figure 9:**
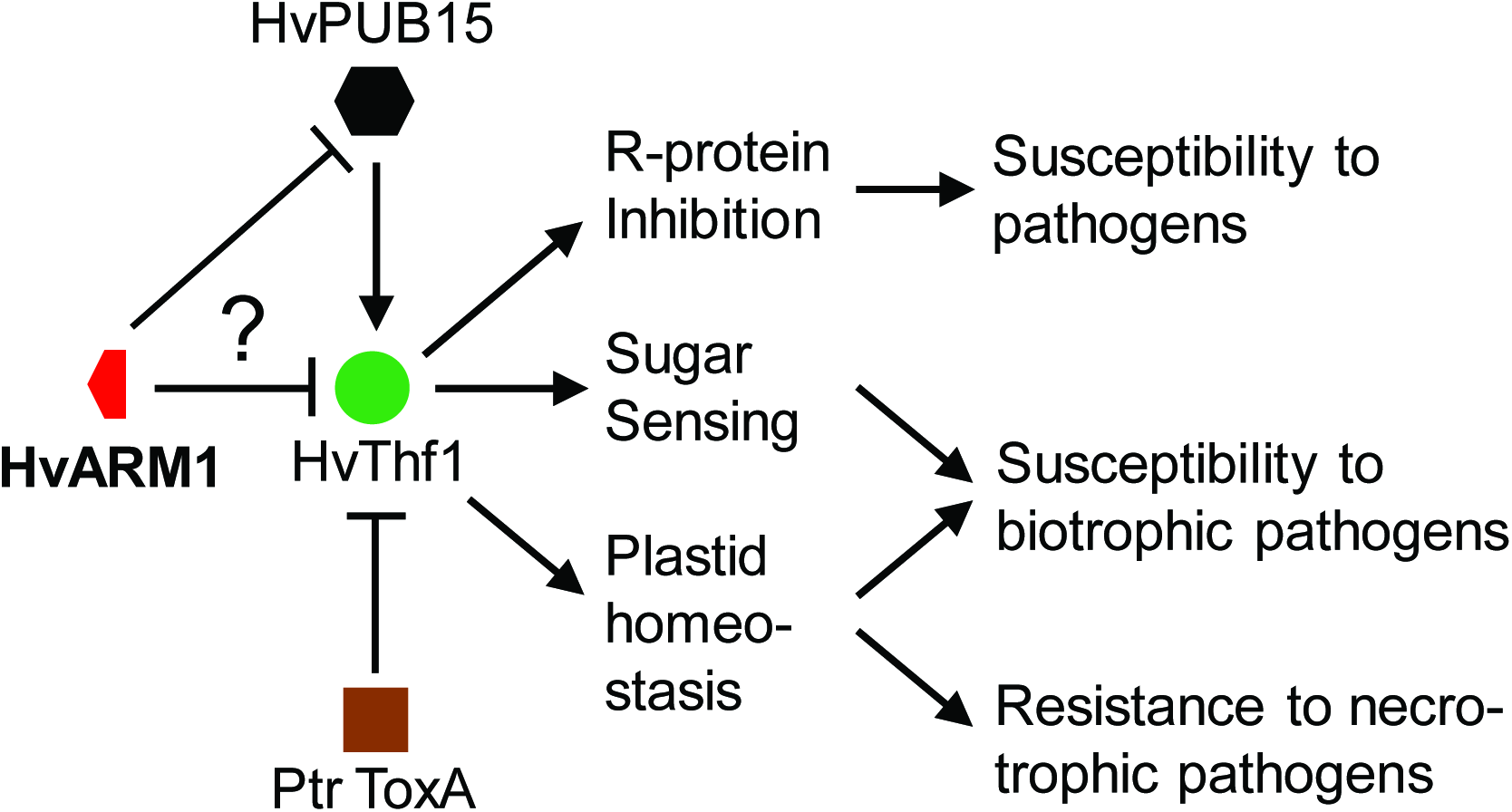
Model of the HvThf1-related functions of HvARM1 and HvPUB15 in powdery-mildew attacked *H. vulgare*. The model is centered on the proposed susceptibility factor HvThf1 for *Bgh*. The Thf1 protein has been proposed as target to the nectotrophic toxin A of *P. tritici-repentis* (Ptr ToxA). The question mark relates to the open possibilities that HvARM1 either directly affects the susceptibility-related function of HvThf1 by binding to it, or that it protects HvPUB15 from putative *Bgh* effector attack.

In summary, our results suggest that *ARM1* was neo-functionalized after a non-tandem, partial gene duplication event of the E3-ligase *PUB15*, which occurred in a common ancestor of the *Triticeae* tribe of grasses and gained a role in broad-spectrum quantitative resistance against *B. graminis* and maybe other pathogenic fungi. At least in barley, the HvARM1-interacting protein and proposed substrate of HvPUB15, the plastid-localized HvThf1, links susceptibility to biotrophic pathogens with plastid functions. Future work for a better understanding of resistance-related ARM1 functions may include the characterization of null-allelic mutants and further allele-mining in barley genetic resources, combined with functional validation and introgression of superior alleles into modern adapted germplasm. The work presented here (Figure 8) also opens up possibilities to search for and functionally address additional partial gene duplicates in crop genomes for a neo-functionalized role in biotic-stress resistance.

## ACKNOWLEDGEMENTS

This work was supported by the German Ministry for Education and Research, grant acronyms GABI-nonhost (to P.S.) and GABI-phenome (to P.S. and J.KU.), by German DFG (ERA-PG project TritNONHOST grant Nr. DFG Schw 848/2-1 to P.S.), by EU FP6 project BIOEXPLOIT (to P.S.) and by EU FP7 project WHEALBI (to P.S. and N.S). We would like to thank Gabi Brantin, Manuela Knauft, Sonja Gentz and Cornelia Marthe for excellent technical assistance.

---

**Table 5.** Transient over-expression of *HvARM1* enhances resistance in *T. aestivum against B. graminis f.sp. tritici*.

| Bombarded gene | Relative SI (log2)<sup>a</sup> | p (t-test)<sup>b</sup> | n<sup>c</sup> |
| --- | --- | --- | --- |
| HvARM1 | -0.37 ± 0.12 | 0.0107 | 16 |
| HvARM1(-ATG)<sup>d</sup> | 0.19 ± 0.13 | 0.2168 | 5 |

<sup>a</sup>Susceptibility index, normalized to the empty-vector control pIPKTA09 and log2-transformed.
<sup>b</sup>One-sample t-test (2-tailed) of log2-transformed relative SI against the hypothetical value “0”.
<sup>c</sup>Number of independent bombardment experiments.
<sup>d</sup>Negative control construct without translation start codon.

## SUPPORTING INFORMATION

**Figure S1:**Alignment of *HvPUB15*and *HvARM1*genomic sequences.

**Figure S2:**Protein alignment of HvARM1 to HvPUB15.

**Figure S3:**Off-target prediction in transgenic barley carrying the RNAi hairpin construct pIPKb009 for silencing of Hv*ARM1*.

**Figure S4:**Expression of Hv*ARM1*endogenous transcripts in powdery mildew-attacked barley epidermis.

**Figure S5:**Localization of YFP-tagged proteins in barley epidermal cells.

**Figure S6:**In vitro ubiquitin ligase activity of HvPUB15.

**Figure S7:**Additional controls for BiFC in *N. bentaminiana*leaves.

**Figure S8:**Quantification of BiFC signals in *N. benaminiana*leaves.

**Figure S9:**Additional, array-based transcript regulation data of selected genes in powdery mildew-attacked barley leaves.

**Figure S10:**Plasmid map of pIPKTA48 for the subcellular localization of N-terminal fusion proteins with YFP.

**Figure S11:**Plasmid map of pIPKTA49 for the subcellular localization of C-terminal fusion proteins with YFP.

**Table S1:**Passport data of the plant accessions used for association genetic analysis.

**Table S2:**SNP calls and derived gene haplotypes based on Exome Capture re-sequencing of the barley genome.

**Table S3:**Interacting candidates from yeast-2-hybrid screening.

**Table S4:**Genome-wide search for pairs of partially duplicated genes.

**Table S5:**Primary signal intensity data from GeneSpring analysis of Agilent 44K Gene Expression array of two selected transcripts shown in supporting Figure S9.

**Table S6:**PCR and TaqMan primers used in the study.

**Methods S1:**More detailed description of Materials and Methods used for the study.

